# Pervasive biases in proxy GWAS based on parental history of Alzheimer’s disease

**DOI:** 10.1101/2023.10.13.562272

**Authors:** Yuchang Wu, Zhongxuan Sun, Qinwen Zheng, Jiacheng Miao, Stephen Dorn, Shubhabrata Mukherjee, Jason M. Fletcher, Qiongshi Lu

## Abstract

Almost every recent Alzheimer’s disease (AD) genome-wide association study (GWAS) has performed meta-analysis to combine studies with clinical diagnosis of AD with studies that use proxy phenotypes based on parental disease history. Here, we report major limitations in current GWAS-by-proxy (GWAX) practices due to uncorrected survival bias and non-random participation of parental illness survey, which cause substantial discrepancies between AD GWAS and GWAX results. We demonstrate that current AD GWAX provide highly misleading genetic correlations between AD risk and higher education which subsequently affects a variety of genetic epidemiologic applications involving AD and cognition. Our study sheds important light on the design and analysis of mid-aged biobank cohorts and underscores the need for caution when interpreting genetic association results based on proxy-reported parental disease history.

## Introduction

Genome-wide association studies (GWAS) have greatly advanced our understanding of the genetic underpinning of complex diseases, unveiling numerous genotype-phenotype associations(*1*). The tremendous success of GWAS in recent years can be attributed, in part, to the emergence of population biobanks, such as the UK Biobank (UKB)(*2*), which provide extensive genotype and phenotype data on large samples. However, a persistent challenge in biobank-based GWAS applications is that these cohorts mainly consist of mid-aged individuals who are too young to have a diagnosis of late-onset diseases. To address this limitation, Liu et al.(*3*) introduced GWAS-by-proxy (GWAX) based on a simple idea—although biobank participants may not have their own diagnosis on late-life disease outcomes, they report their parents’ diagnoses through the family health history survey; they also (indirectly) provide parental genetic data, as their biological child. In their paper, Liu et al. demonstrated the efficacy of GWAX through replicating risk loci identified in case-control GWAS for several diseases including Alzheimer’s disease (AD)(*3*). Since then, GWAX has quickly gained popularity in complex disease genetic research, particularly for neurodegenerative diseases. In fact, GWAX has become so popular in AD genetic studies that every recent AD GWAS performed meta-analysis to combine associations from clinically diagnosed AD cases-controls(*4*) with GWAX proxy associations to boost sample size and statistical power(*5–10*). Further, the largest AD GWAS to date(*9*) has stopped the earlier convention of sharing separate association results for GWAS and GWAX in their study. Instead, only the meta-analyzed association results were made available to the research community.

However, methodological issues in GWAX and the quality of its association results have not been fully investigated. The Liu et al. 2017 paper provided evidence that top genome-wide significant loci yielded similar results in GWAS and GWAX analyses(*3*). Since then, critiques of GWAX have mostly focused on the imprecision and heterogeneity in survey data (i.e., measurement error in parental health history) and their implications in certain genetic applications (e.g., heritability estimation)(*11, 12*). There have been few studies investigating potential systematic biases and methodological limitations in GWAX, particularly regarding the infinitesimal biases that do not appear substantial when focusing on top GWAS loci with large effects but could severely bias applications that involve more complete aggregations of genome-wide association estimates, such as genetic correlation estimation and polygenic risk score (PRS).

In this study, we report evidence of widespread divergent findings between GWAX based on family health history and case-control GWAS for AD(*13*), revealing pervasive biases in current GWAX approaches. We implement GSUB, a GWAS-by-subtraction strategy(*14*), to quantify the biases originating from different sources, revealing that AD GWAX suffers from substantial survival bias from differential parental lifespans, participation bias in the parental health history survey, and reporting bias in the parental health history survey. We demonstrate that almost all existing GWAX approaches produce counter-intuitive and improbable *positive* associations between higher cognition/education and dementia risk. We show that several common genetic epidemiological applications involving AD and cognition yield mixed findings due to this issue in existing AD genetic studies. We also employ a variety of methods to reduce these biases and benchmark their performance. Our findings emphasize an urgent need for caution when interpreting GWAX association results and provide guidance on future study designs involving proxy phenotypes derived from family health history.

## Results

### GWAX replicates top AD risk loci but shows discrepant genetic correlations with other complex traits

To assess the validity of GWAX for AD, we first aimed to replicate the genome-wide significant loci (p ≤ 5e-8) identified in a recent AD case-control study(*13*). We performed GWAX using UKB participants of European descent who reported parental history of AD/dementia (N = 47,993 proxy cases and 315,096 proxy controls; **Methods**). AD GWAX produced similar association results compared to GWAS (**Figure 1A**-**B****, Supplementary Table 1**). Consistent with previous findings(*3*), GWAS and GWAX effect estimates were highly correlated (cor = 0.97 with *APOE* excluded), but we found a substantial attenuation in GWAX effect sizes (regression slope = 0.63). Such an attenuation is not explained by measurement errors in association effects (**Supplementary Figure 1**) but may be explained by winner’s curse: we obtained a regression slope of 1.15 (standard error [se] = 0.20) after correcting for winner’s curse(*15*) (**Methods**). Similar results were found using the top SNPs identified in GWAX (**Supplementary Table 2** and **Supplementary Figure 1**).

**Figure 1.**
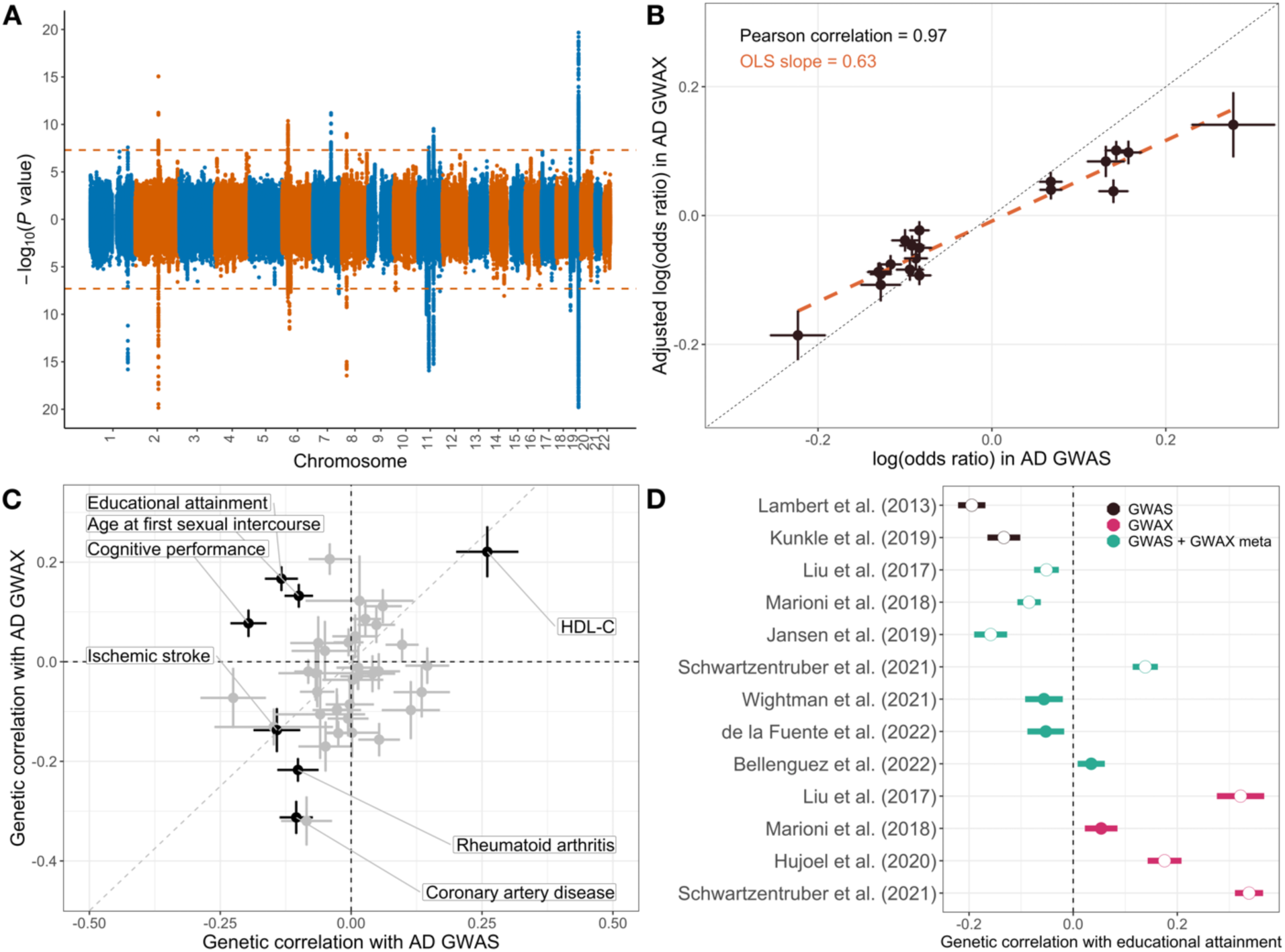
Comparing top association findings and genetic correlation results of AD GWAS and GWAX. (**A**) Manhattan plot for AD GWAX (upper) and GWAS (Kunkle et al. 2019; lower). The y-axis is capped at 20 for better visualization. Horizontal lines mark the genome-wide significance cutoff of 5.0E-8. (**B**) GWAS and GWAX effect size estimates for 20 genome-wide significant SNPs identified in Kunkle et al. 2019. GWAX effect sizes were adjusted using the 0.5 genetic relatedness between parents and children. The *APOE* locus was excluded due to its extreme effect size. (**C**) Genetic correlations of AD GWAX and GWAS with 40 complex traits. Traits with significant correlations (FDR<0.05) with both GWAS and GWAX are highlighted and labeled. HDL-C: high-density lipoprotein cholesterol. (**D**) Genetic correlation of AD and educational attainment based on 10 AD genetic studies published between 2013 and 2022. In (**B**)-(**D**), dots and intervals indicate point estimates and standard errors. Significant results at an FDR cutoff of 0.05 are highlighted with white circles in (**D**).

Discrepancies between GWAX and GWAS became evident in analyses leveraging genome-wide data which include single-nucleotide polymorphisms (SNPs) not reaching Bonferroni-corrected statistical significance. We estimated genetic correlations of AD GWAS and GWAX with 40 complex traits (**Methods**; **Supplementary Tables 3-4**). AD GWAS and GWAX are significantly correlated (cor = 0.63, p = 3.9E-31), but they showed divergent correlations with multiple traits (**Supplementary Figure 2**). For example, total cholesterol and hippocampal volume showed significant genetic correlations with AD GWAS (cor = 0.13 and -0.23, p = 0.01 and 3.1E-4, respectively) but not with GWAX (cor = -0.061 and -0.073, p = 0.23 and 0.23, respectively). Attention-deficit/hyperactivity disorder (ADHD) and coronary artery disease showed substantially stronger correlations with lower AD risk in GWAX (cor = -0.16 and -0.31; p = 2.9E-6 and 3.4E-21) than in GWAS (cor = 0.05 and -0.1; p = 0.18 and 9.5E-4).

Only seven traits had significant correlations with AD in both GWAS and GWAX under a false discovery rate (FDR) cutoff of 0.05, out of which three correlations had flipped directions (**Figure 1C**). In particular, educational attainment (EA), a well-documented negative correlate of AD risk(*16, 17*), showed an expected negative genetic correlation with AD GWAS (cor = -0.13, p = 2.4E-5), but a significant yet positive correlation with AD GWAX (cor = 0.17, p = 1.7E-11). To see if this is a consistent finding across AD studies, we calculated and summarized the genetic correlations between EA and 10 AD studies published between 2013 and 2022(*3, 5–9, 13, 18–20*) (**Supplementary Table 5**). All studies showed a consistent pattern: higher education is correlated with lower AD risk in case-control studies but is correlated with higher AD risk when using family history as the proxy, and results based on GWAS-GWAX meta-analysis fall in-between (**Figure 1D**). The largest AD genetic association study to date(*9*), a meta-analysis of case-control GWAS and family history-based GWAX, showed a positive genetic correlation with EA but did not reach statistical significance (cor = 0.03, p = 0.18).

### GWAX biases risk prediction and causal inference applications involving AD and cognition

Given the concerningly divergent AD-EA genetic correlations based on GWAS and GWAX, we investigated two types of common genetic epidemiologic applications involving AD and cognition. First, we quantified the predictive performance of AD PRS on late-life cognition in the Health and Retirement Study (HRS). We calculated three different PRS from AD GWAS(*13*), GWAX, and a GWAS-GWAX meta-analysis(*9*), and associated these scores with the global cognition composite score while controlling for age, age-squared, education years, year respondent entered study, sex, and top 5 genetic principal components (PCs) (N = 12,018; **Methods**). GWAS-based PRS exhibited a strong association with lower cognition (effect = -0.05, p = 2.5E-11) while GWAX-based PRS did not show significant associations (effect = -0.0017, p = 0.80; **Figure 2A**). PRS based on the Bellenguez et al. 2022 meta-analysis was associated with lower cognition in HRS but showed an attenuated effect size (effect = -0.03, p = 1.2E-7) despite the substantially larger sample size compared to the earlier case-control study. We obtained similar results after removing *APOE* from all PRS (**Supplementary Figure 3**; **Methods**). We also investigated an alternative PRS approach using only variants reaching genome-wide significance in Kunkle et al. 2019 case-control GWAS. The scores based on GWAS and GWAX effects showed very similar performance (**Supplementary Figure 3**), suggesting that biases in PRS analysis were mostly driven by SNPs not reaching statistical significance in AD GWAS.

**Figure 2.**
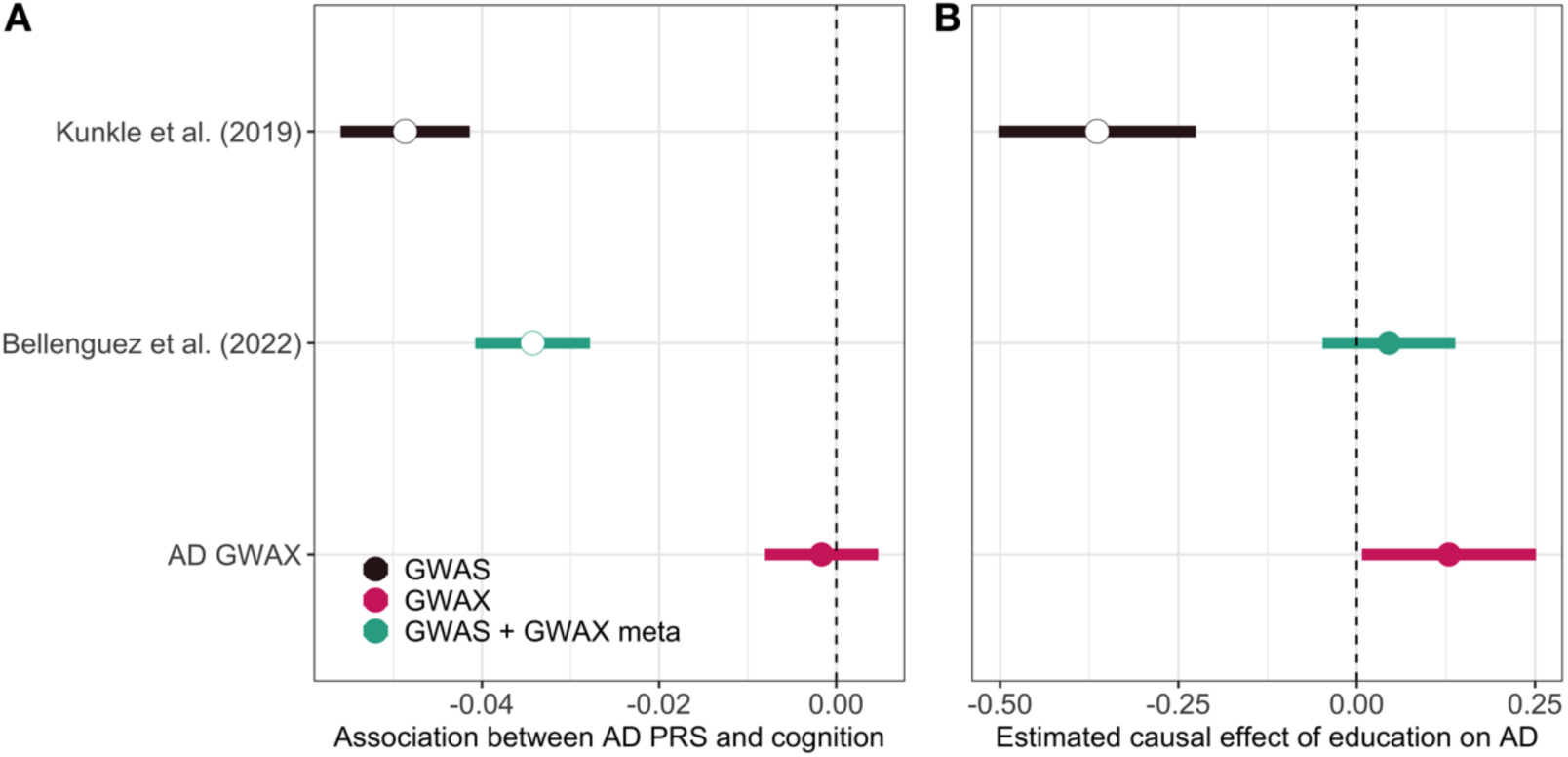
AD GWAX biases risk prediction and causal inference. (**A**) Association of AD PRS and late-life cognition in the HRS cohort. PRS were computed from genome-wide association results using the PRS-CS approach. (**B**) Causal effect of EA on AD risk estimated from Mendelian randomization. For both panels, dots and intervals indicate point estimates and standard errors. Significant results at an FDR cutoff of 0.05 are highlighted with white circles. Data for this plot are in **Supplementary Tables 6-7**.

Education has been hypothesized to have a causal protective effect against AD. Many studies have investigated this hypothesis with mixed results(*16, 17, 21, 22*). Using Mendelian randomization, we estimated the causal effect of EA on AD risk (**Methods**). Once again, we observed inconsistent results between AD GWAS and GWAX (**Figure 2B**). We identified a significant protective effect of EA on lower AD risk using AD case-control GWAS (effect = -0.36; p = 8.6E-3). When GWAX was the outcome study, EA was estimated to increase AD risk although the effect was not statistically significant. A slightly positive but non-significant causal effect of EA on AD risk was also identified using the Bellenguez 2022 meta-analysis. The discrepancies between AD GWAS and GWAX in these analyses underscore the need for a re-evaluation of GWAX applications in AD genetic studies.

### GWAS-by-subtraction identifies potential sources of bias in AD GWAX

Next, we applied a GWAS-by-subtraction approach to separate biases from AD genetic associations in GWAX. This approach assumes that GWAX associations can be explained by real AD signals (i.e., the AD factor F_AD_) and biases (i.e., the non-AD factor F_non_). It quantifies the genetic basis of the non-AD component by regressing out the AD case-control associations from GWAX results (**Methods**; **Figure 3**). GWAS-by-subtraction(*14, 23, 24*) has had several important applications in the literature and is implemented under GenomicSEM(*25*). Our primary analyses using this tool encountered computational singularity issues due to the high genetic correlation between the two input datasets (i.e., AD GWAS and GWAX). Therefore, we applied an alternative strategy to perform GWAS-by-subtraction based on our previous work aimed at decomposing direct and parental genetic effects on children’s outcomes(*26*). This approach produces closed-form estimates for the main parameters of interest, i.e., SNP effects on the non-AD factor F_non_ (**Figure 3**). We have implemented this approach in a software package named GSUB. Compared to GenomicSEM, GSUB produces consistent effect estimates with comparable statistical power. It does not suffer from convergence issues and is computationally much faster (**Supplementary Figure 4; Supplementary Table 8**; **Methods**).

**Figure 3.**
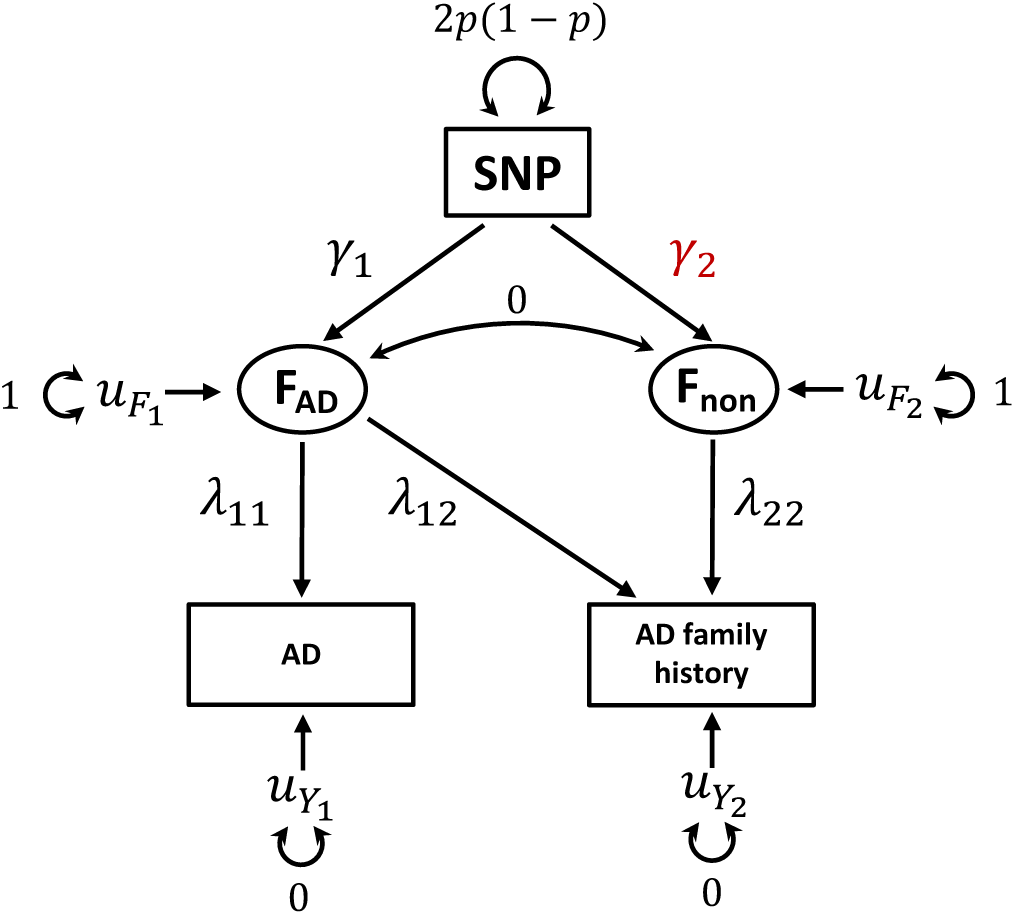
Schematic diagram for GWAS-by-subtraction. The main goal is to estimate genetic associations *γ*_1_ with the non-disease factor F_non_ underlying parental disease history. *γ*_1,2_ and *λ*_11,12,22_ are the parameters that need to be estimated (**Methods**).

To elucidate the mechanisms behind the non-AD (i.e., bias) genetic component underlying AD GWAX, we computed its genetic correlations with 50 complex traits. These include 40 complex traits we have used in previous analysis (**Supplementary Tables 9-10**). In addition, due to EA’s highly divergent genetic correlations with AD GWAS and GWAX (**Figure 1C-D**), we included three additional GWAS measuring different aspects of EA and cognition in the analysis: the direct and indirect (i.e., parental) genetic effects on EA estimated from family-based GWAS(*26*) and the non-cognitive component for EA(*14*). Further, to compare with non-AD dementia, we included GWAS for Parkinson’s disease(*27*), amyotrophic lateral sclerosis(*28*), frontotemporal dementia(*29*), and Lewy body dementia(*30*). Finally, to investigate the effect of non-random participation, we performed GWAS on “do not know parental illness” in UKB (**Supplementary Figures 5; Supplementary Tables 11**), and family medical history awareness and participation of family health history survey (**Supplementary Figures 6-9; Supplementary Tables 12-13**) using data from the AllofUs research program (**Methods**). We included these additional GWAS for genetic correlation estimation, increasing the total number of traits to 50 in this analysis.

**Figure 4** shows significant genetic correlations with the non-AD factor underlying GWAX. 16 traits reached statistical significance at FDR < 0.05. The non-AD GWAX component exhibited substantial correlations with higher EA (cor = 0.26, p = 5.2E-11), indirect (parental) effect on EA (cor = 0.53, p = 4.5E-4), cognition (cor = 0.19, p = 1.4E-4), and the non-cognitive component(*14*) for EA (cor = 0.23, p = 1.4E-6). In addition, we observed negative genetic correlations between the non-AD component with several health outcomes such as major depressive disorder (cor = -0.11, p = 0.012), schizophrenia (cor = -0.13, p = 8.3E-3), coronary artery disease (cor = -0.14, p = 2.8E-3), ADHD (cor = -0.17, p = 0.012), epilepsy (cor = -0.20, p = 7.1E-4), and heart failure (cor = -0.24, p = 4.1E-4), possibly suggesting survival bias in AD GWAX. That is, parents who have AD diagnosis would have to have lived long enough to receive the diagnosis, thus having lower genetic risks for other health issues due to the competing risk. Meanwhile, if some proxy respondents have younger parents who have not reached the age of dementia onset, they will not have lower genetic risks for other outcomes. Therefore, genetic footprints for many health outcomes could partially explain the genetic differences between proxy cases and controls. Indeed, we observed distinct age distributions between GWAX cases and controls (**Supplementary Figure 10**). Compared to proxy AD cases, participants who did not report parental AD history, along with their parents, tended to be younger.

**Figure 4.**
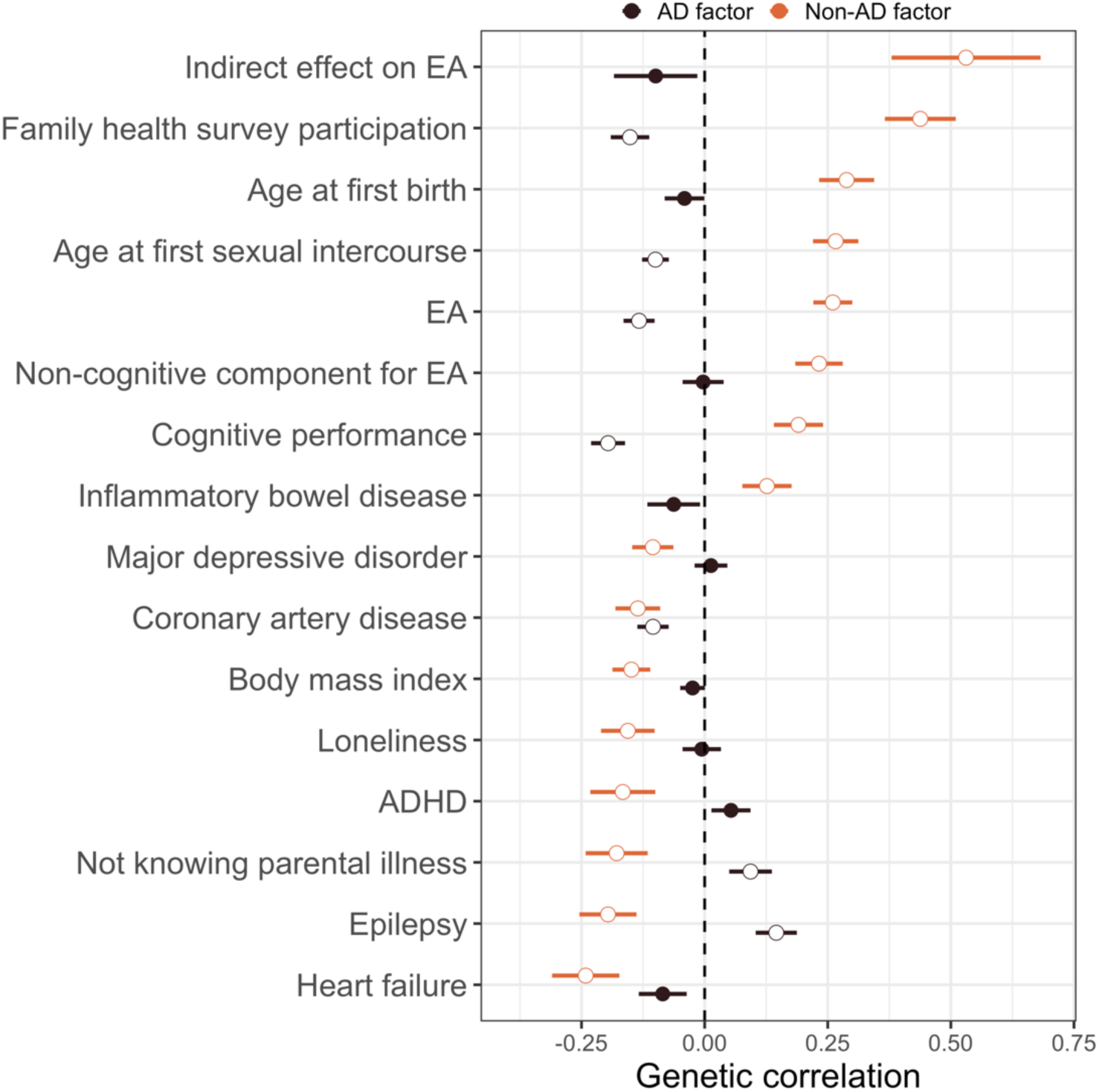
Genetic correlation of the AD and non-AD factors in GWAX with other complex traits. Here, GWAS for the AD factor is Kunkle et al. 2019 case-control GWAS (**Methods**) while genetic associations with the non-AD component were obtained using GWAS-by-subtraction. 16 traits showing significant correlations with the non-AD factor are plotted. Dots and intervals indicate point estimates and standard errors. Significant correlations with FDR < 0.05 are highlighted with white circles. Full genetic correlation results are reported in **Supplementary Tables 3, 9** and **10**.

Since the parental health history survey question in UKB, i.e., “Has/did your father (mother) ever suffer from Alzheimer’s disease/dementia?”, lacks a clear differentiation between AD and other dementia, we examined whether genetic associations for non-AD dementia could explain the biases in AD GWAX. Parkinson’s disease, amyotrophic lateral sclerosis, frontotemporal dementia, and Lewy body dementia all showed null results with genetic correlation estimates close to zero (**Supplementary Table 9**), providing very limited evidence to support this hypothesis.

A recent study(*31*) demonstrated a genetic basis for non-random survey response in UKB. We next investigated whether participation in the family health history survey and systematic misreporting of parental disease status may explain biases in AD GWAX. We found significant genetic correlations between the non-AD component with participation in the family health history survey (cor = 0.44, p = 1.1E-9) and (not) knowing parental illnesses (cor = -0.18, p = 4.6E-3).

### Reducing biases in GWAX

Having identified several potential sources of bias in AD GWAX, we next explored various methods to correct for these biases in GWAX implementation. To reduce survival bias, we applied two approaches in the literature to control for parental age and vital status in the regression (**Supplementary Table 14**). Following Marioni et al. 2018, we excluded parents younger than 65 and added parental age as a covariate in GWAX(*5*); following Jansen et al. 2019, we constructed a continuous GWAX phenotype using parental AD status, their age, and AD prevalence(*6*) (**Methods**). Using AD-EA genetic correlation as a benchmark, both approaches reduced bias in GWAX (**Figure 5**). While the Marioni approach showed a null genetic correlation with EA, the Jansen approach flipped AD-EA genetic correlation from 0.17 to -0.15 which became very close to the genetic correlation based on case-control AD GWAS (cor = -0.13). We also examined the genetic correlation with coronary artery disease as a benchmark for survival bias (**Figure 5**). The Marioni approach substantially reduced the genetic correlation (cor = -0.081, p = 0.01), showing a similar result compared to AD case-control GWAS. The Jansen approach yielded a significant but positive genetic correlation with coronary artery disease (cor = 0.14, p = 1.5E-6).

**Figure 5.**
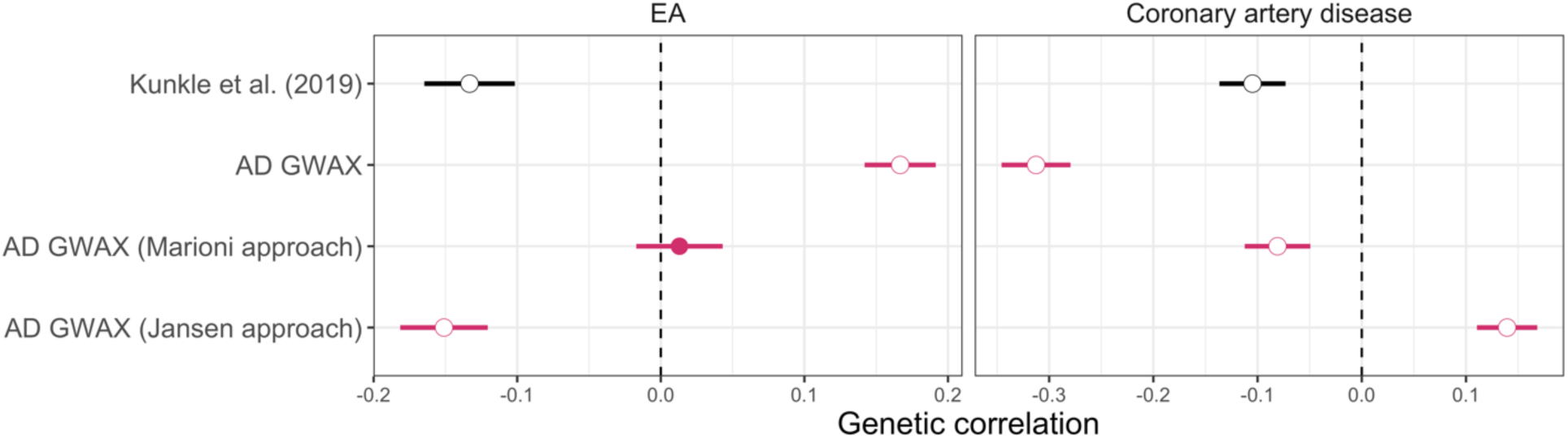
Genetic correlation of UKB AD GWAS and GWAX with educational attainments and coronary artery disease. We included two approaches to correct for survival bias. Following Marioni et al. (2018)’s approach, we require participants parental age (either current age or age at death) to be older than the AD onset age of 65 and including parental age in covariates (**Methods**). Following Jansen et al. (2019), we ran a GWAS on a continuous phenotype which was constructed based on parental AD status, parental age, and AD prevalence to quantify the disease load. Dots and intervals indicate the point estimates and +/- one standard error for the estimate, respectively. Significant results at an FDR cutoff of 0.05 are highlighted with white circles. Data for this plot is in the **Supplementary Tables 14-15**.

To reduce participation bias, we followed Schoeler et al.(*32*) and conducted a weighted GWAS on parental AD status. We trained a LASSO regression model on whether a survey participant reported parental illnesses using a random subset of UKB samples and then performed weighted GWAS on the remaining samples (**Methods**). However, this approach did not improve the genetic correlation estimates with EA (cor = 0.19, se = 0.046, p = 4.3E-5) or coronary artery disease (cor = -0.41, se = 0.078, p = 2.1E-7; **Supplementary Tables 14-15**). We also explored using GWAS-by-subtraction to adjust for participation bias by regressing out the participation GWAS from AD GWAX where the participation GWAS was conducted using AllofUs samples (**Methods**). The residual GWAS showed reduced genetic correlations with EA (cor = 0.11, p = 1.1E-4) and coronary artery disease (cor = -0.18, p = 1.7E-9) but both correlations remained statistically significant (**Supplementary Tables 14-15**).

To address the bias due to systematic over- or under-reporting in the parental health history survey, we explored two different strategies. First, in our default GWAX implementation, we have removed people who reported “do not know” in the parental health history survey, thus already controlling for family health awareness to some extent. To investigate whether this is a reasonable strategy, we implemented another GWAX with people not knowing about parental health added as controls (**Supplementary Table 14**). As expected, after this change we found further inflated genetic correlations between AD and higher EA (cor = 0.21, p = 7.7E-19; **Supplementary Table 14**) and lower risk for coronary artery disease (cor = -0.33, p = 2.5E-26; **Supplementary Table 15**). Additionally, we once again utilized GWAS-by-subtraction, this time regressing out the “do not know parental illness” genetic component from AD GWAX (**Methods**; **Supplementary Figure 11**). We estimated the residual GWAS’ genetic correlations with EA and coronary artery disease and obtained substantially reduced yet still significant correlations (cor = 0.07, p = 0.027 with EA; cor = -0.26, p = 1.3E-13 with coronary artery disease; **Supplementary Tables 14-15**).

Finally, we note that although both the Marioni and Jansen approaches were primarily designed for reducing survival bias alone, they also removed some reporting bias. After correction, AD GWAX had null genetic correlations with “do not know parental illness” (cor = -0.03 and 0.087, p = 0.54 and 0.06 for Marioni and Jansen approaches, respectively; **Supplementary Tables 16-17**).

### Meta-analysis of GWAS and GWAX associations

AD GWAX is often meta-analyzed with clinically diagnosed case-control GWAS summary statistics to boost statistical power. Next, we investigate whether accounting for heterogeneity when meta-analyzing GWAS and GWAX could reduce biases in the combined association results. We explored two approaches: METAL(*33*) is a common approach for meta-analysis and GenomicSEM was recently proposed as an alternative strategy that can account for measurement error and phenotype heterogeneity in GWAX-GWAS meta-analysis(*12, 20*). **Figure 6** illustrates the genetic correlations of the meta-analyzed outcomes based on two meta-analytic approaches with EA and coronary artery disease. Compared to results in **Figure 5**, meta-analyzing GWAX with GWAS produces genetic correlations somewhere in-between those given by GWAX and GWAS. It was also clear that meta-analysis alone cannot sufficiently remove all the bias. The two meta-analytic methods produced mostly comparable results, highlighting the importance of reducing biases in GWAX analysis instead of relying solely on *post hoc* bias reduction during meta-analysis.

**Figure 6.**
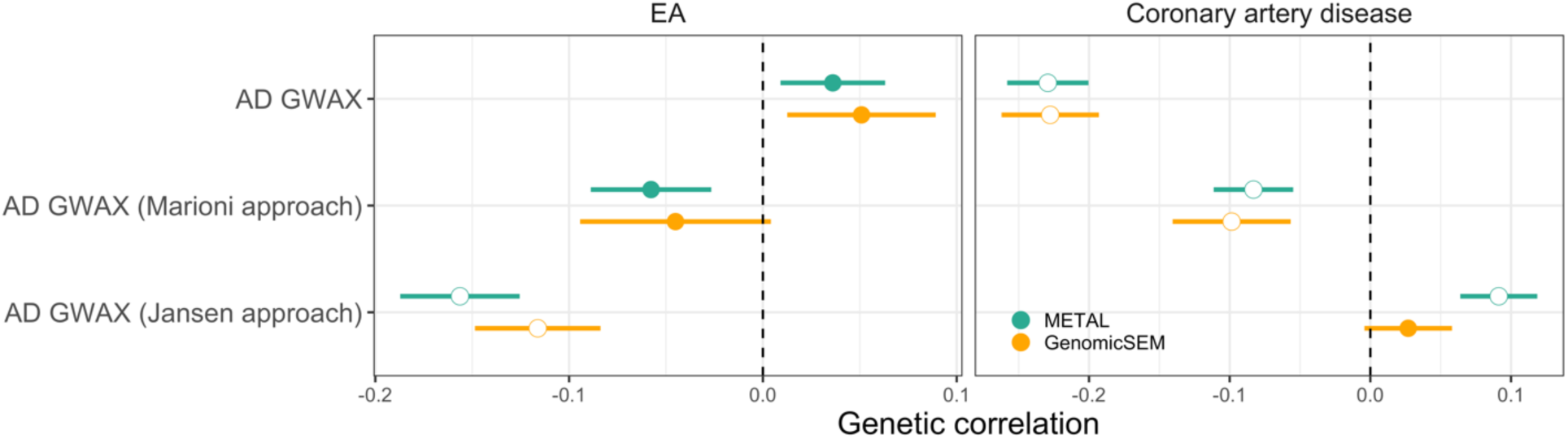
Genetic correlation of meta-analyzed AD with EA and coronary artery disease. We show meta-analysis results based on GWAS from Kunkle et al. (2019) and three sets of AD GWAX. Results based on two meta-analysis approaches, i.e., METAL(*33*) and GenomicSEM(*25*), are also compared. We used METAL to combine GWAX based on parental AD history with GWAS associations. Since GenomicSEM requires at least three studies as input, we meta-analyzed GWAS, paternal GWAX, and maternal GWAX. Data for this plot is in **Supplementary Tables 14-15**.

## Discussion

In recent years, GWAX has emerged as a crucial study design for complex trait genetics in general and AD genetic research in particular, gaining popularity due to its ability to leverage mid-aged population data to study late-onset health outcomes. The validity of GWAX was supported by two types of evidence in the literature: similar effect size estimates for top SNP findings and high genetic correlation between GWAS and GWAX based on genome-wide data. Some critiques have been raised concerning the GWAX design, mostly focusing on measurement errors in family health history survey. Escott-Price and Hardy(*11*) argued that parental AD cases inferred from vaguely defined surveys may encompass both AD and non-AD dementia cases, each with distinct genetic underpinnings, which would attenuate genuine genetic associations for AD. More recently, Grotzinger et al. demonstrated that naively combining GWAS and GWAX without accounting for heterogeneity among the associations will lead to substantial downward bias in heritability estimation(*12*). Despite these critiques, GWAX has become an integral component of every recent AD GWAS(*5–10*), raising concerns about quality of reported associations and prospect of follow-up studies based on GWAS findings.

In this paper, we revealed pervasive and systematic biases in AD GWAX associations. In particular, AD GWAX yielded an unexpected positive genetic correlation with EA, and such biases are present in almost all published AD GWAS that included proxy AD cases. The significance of this issue is two-fold. First, the biases identified in our analyses are not just speculations of some negligible issue in empirical applications. We demonstrated substantial divergence of AD GWAS and GWAX in some aspects due to these biases. Second, an important social factor at the center of many of these biases is education – it is known to associate with longevity, parent-child relationship, and general health awareness(*34*). But because cognition is such a crucial marker for AD and is commonly used in dementia research, biases caused by education/cognition become particularly important in AD genetics research and may give misleading results if not handled properly, complicating diagnosis, treatment, and the design and testing of new drugs. We investigated two types of analyses that are frequently done in genetic epidemiology studies of AD: predicting late-life cognition using AD PRS and estimating causal (protective) effect of education on AD using Mendelian randomization. Indeed, both analyses are substantially influenced by biases in GWAX.

Despite the strong evidence for bias, the source of such bias was not clearly understood. In addition to genuine AD associations, we hypothesized that there could be at least three types of mechanisms contributing to bias in GWAX findings. First, only people with parents who lived long enough can report parental AD diagnosis. Without adjusting for such survival bias, we expect to see spurious negative genetic correlations between AD GWAX and other health outcomes. That is, genetic variants that are protective for other diseases will appear to increase the risk of AD because they increase longevity. Second, people who are more aware of their parents’ health are more likely to report parental AD diagnosis. This could be affected by people’s general awareness of health issues, but may also be explained by people’s relationship with their parents, whether they grew up in single-parent families, parents’ socioeconomic status, and other complex socio-environmental factors. Third, parental AD cases reported in the UKB parental health survey may include non-AD dementia cases. Therefore, we expect genetic associations with other types of dementia to explain some differences between AD GWAS and GWAX. Using an innovative GWAS-by-subtraction strategy(*14*) (with our novel closed-form implementation), we quantified genetic effects underlying AD GWAX that are not explained by genuine AD associations. We found substantial evidence for survival bias, supported by negative genetic correlations of the non-AD (bias) component with many health outcomes. We also found genetic correlations with survey participation and awareness of parental health history, which suggest non-random participation and reporting in UKB survey as a possible source of bias. We did not find evidence for other dementia associations in AD GWAX, although this is possibly explained by the lower statistical power in current non-AD dementia studies.

We also investigated several approaches to reduce biases in AD GWAX. We demonstrated that controlling for parental age and vital status could effectively reduce survival bias. In particular, the approach which creates a continuous disease risk phenotype based on parental age(*6*) produced education genetic correlation results comparable to AD case-control GWAS. However, one potential limitation of this approach is that it does not produce SNP effect sizes on a similar scale to case-control studies, which creates challenges in the interpretation and some applications requiring effect sizes. While weighted least squares is a common approach to account for non-random study participation(*32*), it did not give promising results in our analysis. Excluding individuals who do not know about parental health from the analysis and residualizing GWAX on genetic associations with parental health awareness both reduced the spurious AD-EA genetic correlation. We note that the approaches designed to remove survival bias also reduced some participation and reporting biases, suggesting entangled mechanisms behind these possibly over-simplified labels for different sources of bias. This provides a potential one-stop solution to multiple sources of bias but its effectiveness still remains to be carefully investigated in the future. We also note that these biases could not be mitigated by a simple meta-analysis with AD case-control GWAS, further highlighting the importance of improving the quality of GWAX analysis. Finally, besides the issues we have detailed in this study, many association mapping approaches being used in GWAX studies appear statistically poorly justified. For example, AD GWAX sometimes combine clinically diagnosed cases and proxy cases together in logistic regression without properly scaling the SNP effect size according to the proxy case-control design(*7, 9*). Some studies combine both sibling and parental proxy cases(*3, 7, 9*) which could introduce additional survival bias and other complications. Some other studies meta-analyze GWAX associations based on maternal and paternal AD histories without accounting for the sample overlap between them(*5*). There is an urgent need to improve the general statistical methodology for handling family history outcomes in genetic association studies.

Our study has several limitations. First, we treated the AD case-control GWAS as the gold standard throughout the paper, but it remains plausible that some issues could also affect the analysis based on AD clinical diagnosis. For example, the significant genetic correlation between Kunkle et al. 2019 GWAS and lower risk for coronary artery disease (**Figure 5**) is suggestive of uncorrected survival bias in AD GWAS. In fact, GWAX following the Jansen approach showed a positive genetic correlation with coronary artery disease. It is unclear if this is caused by limitations in the bias-removal approach or correctly recovering shared genetics between AD and cardiovascular disease risk(*35, 36*). Second, it is unclear what metrics should be used to benchmark the performance of GWAX. In this study, we used the genetic correlations of AD with EA and coronary artery disease to quantify the effectiveness of bias-reduction approaches. But fully addressing issues in GWAX would require replication and functional validation of findings. Third, the non-presentiveness of UKB participants is well-documented(*37–39*), but it has been suggested that some sampling issues in UKB are not observed in other cohorts(*38*). Additionally, we only focused on individuals of European descent in our analysis. It is an important future direction to investigate how these issues generalize to other ancestries and cohorts.

Taken together, our findings bear significant implications for the field, as they uncover an urgent, ubiquitous, yet understudied problem hidden in plain sight. Given its popularity and the potential of creating misleading results, it is of great urgency to reassess the statistical foundation of GWAX. We urge the research community to critically reconsider their utilizations of family history-based proxy phenotypes and adopt a more cautious and rigorous approach when drawing conclusions based on GWAX findings. An immediate remedy for all future studies is to release separate GWAS and GWAX summary statistics for research use, although fully addressing these issues will most likely require tremendous efforts in results validation and development of novel statistical methodologies.

## Methods

### GWAS analysis in UKB

We conducted GWAS in UKB for parental AD history and parental illness awareness using Regenie(*40*) while controlling sex, year of birth, and genotyping array (data field 22000 in UKB) as fixed effect covariates. Population stratification was accounted for in the ridge regression step of Regenie which is similar to a linear mixed model approach without having to compute the genetic relatedness matrix. We excluded participants with conflicting genetically inferred (data field 22001) and self-reported sex (data field 31), those who withdrew from the study, and those that are recommended to be excluded by UKB (data field 22010). Individuals of European ancestry were identified from principal component analysis (data field 22006). We kept only the SNPs with a missing call rate ≤ 0.01, minor allele frequency ≥0.01, and Hardy–Weinberg equilibrium test P value ≥1E-6.

Parental AD history (i.e., the outcome in AD GWAX) was derived from survey responses to questions regarding the “illnesses of father” (data-field 20107) and “illness of mother” (data-field 20110). Responses options include “Do not know”, “Prefer not to answer”, “None of the above”, or one of the twelve diseases including “Alzheimer’s diseases/dementia”. Participants were coded as proxy cases if either parent had AD, and as controls if both parents were not affected by AD. Samples were removed from analysis if otherwise. Additionally, participants self-identified as adopted (data-field 1767) were excluded from the study. We identified 47,993 proxy cases and 315,096 controls in the parental AD GWAX.

The parental illness awareness phenotype was derived from UKB data fields 20107 and 20110. Cases were those who selected “Do not know (group 1)” or “Do not know (group 2)” for either father or mother’s illnesses. Controls were those who selected “None of the above” or any disease in both groups for both father and mother’s illnesses. Others were excluded from the analysis. 59,471 cases and 339,170 controls were identified.

### GWAS analysis in AllofUs

The AllofUs research program is a nationally representative cohort in the US with a goal of recruiting 1 million participants. We conducted GWAS using AllofUs samples for two phenotypes: participation of the family health history survey and family medical history awareness. The family health history survey is an optional module and only a subset of AllofUs samples participated in this module. We determined survey participation status by checking whether an individual answered the first question in this module, which reads, “How much do you know about illnesses or health problems for your parents, grandparents, brothers, sisters, and/or children?” This question has four possible response options: “none at all”, “some”, “a lot”, and “skip”. The GWAS on family medical history awareness was based on the answers to this question. We coded the responses as follows, “none” as 0, “some” as 1, and “a lot” as 2. Individuals selecting “skip” were excluded from the analysis.

For both GWAS in AllofUs, we used independent samples of European descent and adjusted for biological sex, standardized age, square of the standardized age, and top 16 genetic PCs. GWAS was performed using Hail on version 7 of the whole genome sequencing data. Genetic ancestry inferred from PCs and genetic relatedness between participants were provided in AllofUs. Samples flagged as outliers were excluded from analysis. We kept only the SNPs with a missing call rate ≤ 0.01, minor allele frequency

≥ 0.01, and Hardy–Weinberg equilibrium test P value ≥ 1E-6. Sample size for the GWAS on family medical history awareness is 77,579. There were 78,027 cases (participants) and 47,519 controls (non-participants) in the GWAS on participation of the family medical history survey.

### Measurement error and winner’s curse correction

We used Deming regression implemented in R package “mcr” to correct for measurement errors in SNP effect estimates. We used the “mcreg()” function and specified the ratio of the error variances to be 42,706/41,679 where 42,706 is the effective sample size (sum of all the effective sample sizes from all contributing cohorts) for Kunkle et al. (2019) AD GWAS, and 41,649 is the effective sample size for the AD GWAX we performed in UKB. We used R package “WinCurse” to correct for winner’s curse in the Kunkle et al. (2019) GWAS. The adjusted SNP effect size followed formulas in Turley et al.(*41*)

### Heritability and genetic correlation estimation

We used GNOVA(*42*) to estimate genetic correlations. We corrected GWAS sample overlap in GNOVA if bivariate LDSC(*43*) outputs an intercept significantly different from zero at P < 0.05. We used LDSC to estimate heritability.

### PRS regression analysis

We evaluated the performance of AD PRS in HRS. The HRS is a nationally representative longitudinal biennial panel consisting of around 42,000 Americans from 26,000 households since 1992. A global cognition composite score was derived from a 27-point scale that includes: 1) an immediate and delayed 10-noun free recall test to measure memory (0 to 20 points); 2) a serial sevens subtraction test to measure working memory (0 to 5 points); and 3) a counting backwards test to measure speed of mental processing (0 to 2 points). There are 10 waves of data available: once every 2 years from 2000 to 2018.

We obtained imputed genetic data from a subset of around 15,000 participants who had their genetic information collected between 2006 and 2012 (NIAGADS accession number NG00119.v1). PRS were calculated using two different approaches: PRS-CS(*44*) and clumped significant SNPs in Kunkle et al. 2019 GWAS(*13*). Only overlapping SNPs that exist in all GWAS summary statistics as well as HRS genotype data were used. We used the PRS-CS-auto implementation to estimate SNP posterior effect sizes from genome-wide summary statistics. The second PRS approach weighted allele counts with effect sizes obtained from GWAS summary statistics and only included independent SNPs reaching genome-wide significance in Kunkle et al. 2019 GWAS. Clumping was executed using PLINK1.9, with clumping parameters set at r2 of 0.1 and kb of 1000. We also generated an additional set of PRS excluding the *APOE* region by removing all SNPs in the region (chr19: 45,116,911-46,318,605; GRCh37).

To analyze longitudinal cognition data in HRS, we used random intercepts in linear mixed model to account for within-sample (repeated measures) and within-family (related samples) correlations. The regression analyses were performed using the “lme4” package in R, where we regressed cognitive scores against PRS while controlling for age, age-squared, education years, year respondent entered study, sex, and top 5 genetic PCs. Individual and family IDs were coded as random effects. Only HRS participants of European descent were included in the analysis with a total sample size of 12,018.

### Causal effect estimation

Mendelian randomization was conducted using the “TwoSampleMR” package in R (*45*). To infer the causal effect of EA on AD, we first clumped EA GWAS summary statistics with r2 = 0.001, kb = 10000 in PLINK1.9, then we selected only SNPs reaching genome-wide significance (p < 5E-8) as instruments. The “mr” function was used to estimate causal effects based on the inverse variance weighted approach.

### GSUB: a new implementation for GWAS-by-subtraction

Consider the GWAS-by-subtraction model shown in **Figure 3** where we aim to subtract genuine AD associations from GWAX associations based on AD family history (i.e., decomposing GWAX into AD and non-AD components). There are five parameters to estimate (i.e., *λ*_11,12,22_ and *γ*_1,2_) and the main parameter of interest (i.e., the SNP effect *γ*_2_ on the non-AD component) is highlighted in red. First, we can write the expressions for AD and AD family history phenotypes in the liability scale:

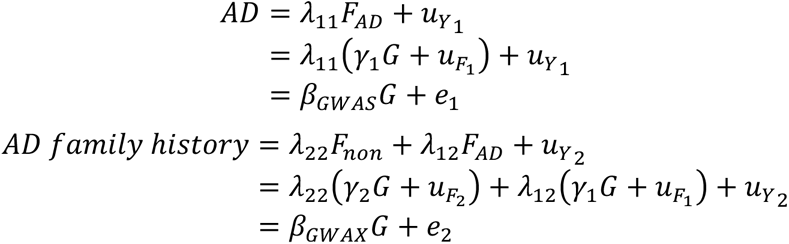

The variances and covariances of the genetic components of the two phenotypes are:

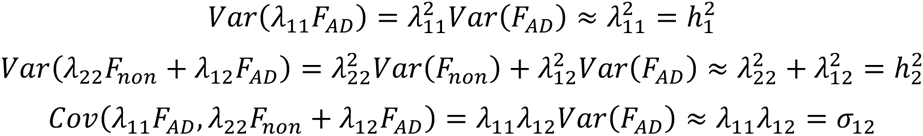

Here, *G* is the SNP allele count, *F_AD_* and *F_non_* are the two latent factors (with variance of 1) underlying AD and AD family history. *β_GWAS_* and *β_GWAS_* are the SNP effect sizes in GWAS and GWAX, respectively. *u* and *e* are residuals. From the first two equations, we have

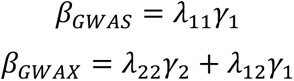

Based on this, we obtain the expressions for *γ*_1_ and *γ*_2_:

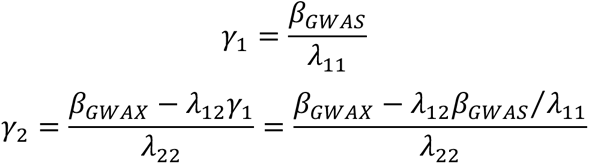

From the third to fifth equations, we could solve for the 3 loading factors:

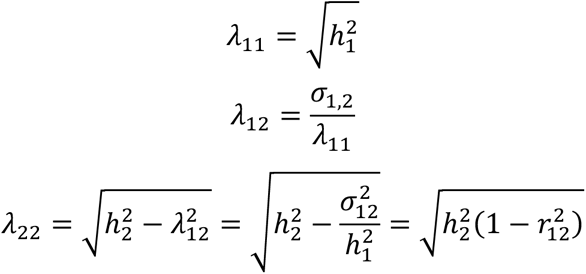

In order to estimate the five parameters, we can plug in the SNP effect size estimates and their standard errors from the summary statistics, the LDSC heritability estimates and genetic covariance between the two traits:

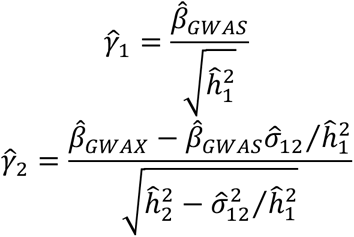

The standard error for *γ̂*_1_ can be approximated by

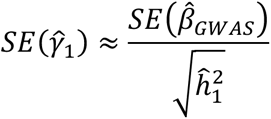

We note that based on this model setting, GWAS for the AD factor is essentially very similar to the input AD case-control GWAS.

To obtain the standard errors for *γ̂*_2_, we need the covariance between *β̂*_1_ and *β̂*_2_. When there are sample overlaps between GWAS and GWAX, their covariance can be estimated using the intercept from the bivariate LDSC:

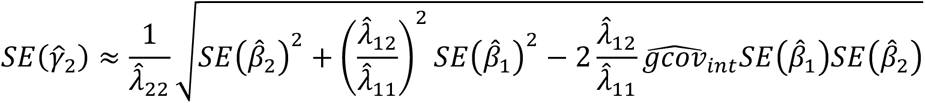

where 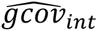 is the bivariate LDSC intercept.

We note that similar derivations for the point estimate of *γ̂*_1_ have been previously shown in the supplementary note of Demange et al. (2021). Here, we provide details for the standard error estimation and have implemented the approach as an open-source software.

### Simulations

We conducted simulations to compare our analytical approach for GWAS-by-subtraction with GenomicSEM. We used HapMap 3 SNP genotype data (853,041 SNPs) from independent UKB samples of European descent. We performed simulations for both quantitative traits (N = 200,000 and 100,000) and binary traits (N = 100,000; case proportion = 20% and 10%). Each setting was repeated 100 times.

Following **Figure 3**, we first simulated SNP effect sizes on each latent factor from a normal distribution with mean 0 and variance 1/M, where M is the number of causal SNPs. The effect sizes were then transformed by dividing 2*p*(1 − *p*), where *p* is the minor allele frequency of each SNP. The latent factors F_1_ and F_2_ were computed as 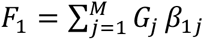 and 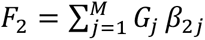, respectively, where *G*_0_ is the allele count (i.e., 0, 1, or 2) for the j^th^ SNP. Then, we calculated the observed continuous trait or disease liabilities as follows.

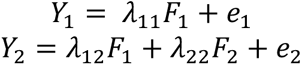

For binary trait, we set samples at the top 10% or 20% disease liability as cases and others as controls. In each repeat, we randomly selected 10,000 causal SNPs for each latent factor. We set 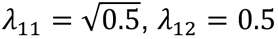, and 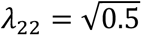.

After simulating phenotype values, we performed GWAS using Plink2.0 for each phenotype. Then, we applied GWAS-by-subtraction using both GenomicSEM and GSUB to compare the type-I error and power. Due to the computational burden of GenomicSEM, we randomly selecting 10,000 null SNPs for type-I error calculation in each repeat. Type-I error (and power) were calculated as the percentage of null (and causal) SNPs with P values < 0.05.

### Approaches for bias reduction in GWAX

We explored several strategies to reduce biases in GWAX. To address survival bias, we implemented two approaches. Following Marioni et al. 2018, we required both parents to be older than 65 which was determined by either current age (data-fields 2946 and 1845) or age at death (data-fields 1807 and 3526). We also included parental age (either current age or age at death) as a fixed-effect covariate. There were 36,309 cases and 199,969 controls in this GWAX. Following Jansen et al. 2019, we created a continuous “disease load” based on parental AD status, parental age, and AD prevalence in the population: each affected parent contributed 1 while each unaffected parent contributed *min*{(100 − *age*)⁄100, 0.32} to the disease load phenotype, where 0.32 is the population prevalence of AD. Those with unknown parental AD status or parental age were excluded from the analysis giving a sample size of 355,501. We performed GWAS on this continuous outcome while controlling for sex, genotyping array, year of birth, and assessment center (data field 54).

We used a weighted GWAS approach to account for non-random survey participation. Following Schoeler et al.(*32*), we used 14 variables to train a participation prediction model. These variables comprise five continuous ones: age, body mass index, weight, height, and the age at which full-time education was completed, and nine categorical variables: household size (1-7 or more individuals), sex (male or female), alcohol consumption frequency (never to daily), smoking habits (never, previous, or current smoker), employment status (employed, economically inactive, retired, or unemployed), income brackets (from <18K to >100k), obesity classification (underweight, healthy weight, overweight, or obese), general health status (poor, fair, or good), and urbanization level (from village/hamlet to urban). We identified 28,179 independent non-reporting individuals (i.e., the non-adopted European ancestry samples that were not included in AD GWAX) for parental AD history. We then randomly sampled the same numbers of individuals who reported parental illnesses to match with these non-reporting individuals. We used LASSO regression in “glmnet” package in R to predict the reporting of parental illnesses. The model included all main effects and two-way interaction terms, with the shrinkage parameter lambda being determined via 5-fold cross-validation. We then conducted weighted GWAS on parental AD history with remaining samples using weighted least squares in R. We used the Huber-White estimator for the variance of the estimates implemented in the “sandwich” package in R. The sampling weights were calculated as *w* = (1 − *p*)⁄*p*, where *p* represents the probability of reporting, predicted through the trained LASSO model. GWAS covariates included sex, year of birth, year of birth squared, genotyping array, and the top 20 PCs. In addition, we also explored using GWAS-by-subtraction to remove participation bias, where we regressed out the participation GWAS from the UKB AD GWAX (**Supplementary Table 14**). The participation GWAS was conducted using the AllofUs samples which we have described in detail.

We applied two approaches to adjust for the reporting bias. The first approach is to include those who selected “Do not know” when answering illnesses of father or illnesses of mother in the analysis as controls, and then repeated the AD GWAX (N = 47,993 cases and 349,165 controls). The second approach is to apply GWAS-by-subtraction to regress out the GWAS on parental illness awareness from AD GWAX.

## URLs

GSUB: https://github.com/qlu-lab/GSUB

R package “mcr”: https://cran.r-project.org/web/packages/mcr/index.html

R package “WinCurse”: https://github.com/zrmacc/WinCurse/tree/master

R package “TwoSampleMR”: https://mrcieu.github.io/TwoSampleMR/

R package “sandwich”: https://cran.r-project.org/web/packages/sandwich/index.html

Regenie: https://github.com/rgcgithub/regenie

GNOVA: https://github.com/qlu-lab/GNOVA-2.0

LDSC: https://github.com/bulik/ldsc

## Data and code availability

Summary statistics for the AD GWAX are available at http://qlu-lab.org/data.html. GSUB software is freely available at https://github.com/qlu-lab/GSUB.

## Acknowledgements

The authors gratefully acknowledge research support from the National Institute on Aging (NIA) grant R21 AG085162, NIA Center Grant P30 AG017266, and support from the University of Wisconsin-Madison Office of the Chancellor and the Vice Chancellor for Research and Graduate Education with funding from the Wisconsin Alumni Research Foundation (WARF). The All of Us Research Program is supported by the National Institutes of Health, Office of the Director: Regional Medical Centers: 1 OT2 OD026549; 1 OT2 OD026554; 1 OT2 OD026557; 1 OT2 OD026556; 1 OT2 OD026550; 1 OT2 OD 026552; 1 OT2 OD026553; 1 OT2 OD026548; 1 OT2 OD026551; 1 OT2 OD026555; IAA #: AOD 16037; Federally Qualified Health Centers: HHSN 263201600085U; Data and Research Center: 5 U2C OD023196; Biobank: 1 U24 OD023121; The Participant Center: U24 OD023176; Participant Technology Systems Center: 1 U24 OD023163; Communications and Engagement: 3 OT2 OD023205; 3 OT2 OD023206; and Community Partners: 1 OT2 OD025277; 3 OT2 OD025315; 1 OT2 OD025337; 1 OT2 OD025276. In addition, the All of Us Research Program would not be possible without the partnership of its participants. The Health and Retirement Study (HRS) genetic data were accessed through NIAGADS with accession number NG00119.v1. These data were collected with financial support from the National Institute of Health’s (NIH) Director’s Opportunity for Research awards using American Reinvestment and Recovery Act funds (RC2 AG036495-01, RC4 AG039029-01). With these funds, the HRS has genotyped almost 20,000 respondents who provided DNA samples and signed consent forms in 2006–2012. The HRS data were produced and distributed by the University of Michigan under the directorship of David R. Weir, with funding from the National Institute on Aging (grant number NIA U01AG009470), Ann Arbor, MI. This research has been conducted using the UK Biobank Resource under Application Number 42148. We thank members of the Social Genomics Working Group at University of Wisconsin-Madison for helpful comments.

## Author contribution

Q.L. conceived and designed the study.

Y.W. and Z.S. performed the analyses.

Y.W. and Q.L. wrote the manuscript.

S.D. implemented the software package for GWAS-by-subtraction.

Q.Z performed the GWAS on parental illness awareness in UKB.

J.M assisted with the mathematical derivations for GWAS-by-subtraction.

S.M. advised on the genetics of AD and cognition.

J.F. advised on the HRS cohort and social science issues.

All authors revised and approved the manuscript.

## References

1. A. Abdellaoui, L. Yengo, K. J. Verweij, P. M. Visscher, 15 years of GWAS discovery: Realizing the promise. The American Journal of Human Genetics, (2023).

2. C. Bycroft et al., The UK Biobank resource with deep phenotyping and genomic data. Nature 562, 203–209 (2018).

3. J. Z. Liu, Y. Erlich, J. K. Pickrell, Case–control association mapping by proxy using family history of disease. Nature Genetics 49, 325–331 (2017).

4. G. McKhann et al., Clinical diagnosis of Alzheimer’s disease: Report of the NINCDS-ADRDA Work Group under the auspices of Department of Health and Human Services Task Force on Alzheimer’s Disease. Neurology 34, 939 (1984).

5. R. E. Marioni et al., GWAS on family history of Alzheimer’s disease. Translational Psychiatry 8, 99 (2018).

6. I. E. Jansen et al., Genome-wide meta-analysis identifies new loci and functional pathways influencing Alzheimer’s disease risk. Nature Genetics 51, 404–413 (2019).

7. J. Schwartzentruber et al., Genome-wide meta-analysis, fine-mapping and integrative prioritization implicate new Alzheimer’s disease risk genes. Nature Genetics, (2021).

8. D. P. Wightman et al., A genome-wide association study with 1,126,563 individuals identifies new risk loci for Alzheimer’s disease. Nature Genetics 53, 1276–1282 (2021).

9. C. Bellenguez et al., New insights into the genetic etiology of Alzheimer’s disease and related dementias. Nature Genetics 54, 412–436 (2022).

10. R. Sherva et al., African ancestry GWAS of dementia in a large military cohort identifies significant risk loci. Mol Psychiatry, (2022).

11. V. A.-O. Escott-Price, J. Hardy, Genome-wide association studies for Alzheimer’s disease: bigger is not always better. Brain Communications 4, (2022).

12. A. D. Grotzinger, J. d. l. Fuente, F. Privé, M. G. Nivard, E. M. Tucker-Drob, Pervasive Downward Bias in Estimates of Liability-Scale Heritability in Genome-wide Association Study Meta-analysis: A Simple Solution. Biological Psychiatry, (2022).

13. B. W. Kunkle et al., Genetic meta-analysis of diagnosed Alzheimer’s disease identifies new risk loci and implicates Aβ, tau, immunity and lipid processing. Nature Genetics 51, 414–430 (2019).

14. P. A. Demange et al., Investigating the genetic architecture of noncognitive skills using GWAS-by-subtraction. Nature Genetics 53, 35–44 (2021).

15. C. A. Rietveld et al., Common genetic variants associated with cognitive performance identified using the proxy-phenotype method. PNAS 111, 13790–13794 (2014).

16. S. C. Larsson et al., Modifiable pathways in Alzheimer’s disease: Mendelian randomisation analysis. BMJ 359, j5375 (2017).

17. S. J. Andrews et al., Causal Associations Between Modifiable Risk Factors and the Alzheimer’s Phenome. Annals of Neurology 89, 54–65 (2021).

18. J.-C. Lambert et al., Meta-analysis of 74,046 individuals identifies 11 new susceptibility loci for Alzheimer’s disease. Nature Genetics 45, 1452–1458 (2013).

19. M. L. A. Hujoel, S. Gazal, P.-R. Loh, N. Patterson, A. L. Price, Liability threshold modeling of case–control status and family history of disease increases association power. Nature Genetics 52, 541–547 (2020).

20. J. de la Fuente, A. D. Grotzinger, R. E. Marioni, M. G. Nivard, E. M. Tucker-Drob, Integrated analysis of direct and proxy genome wide association studies highlights polygenicity of Alzheimer’s disease outside of the APOE region. PLOS Genetics 18, e1010208 (2022).

21. H. Liu et al., Mendelian randomization highlights significant difference and genetic heterogeneity in clinically diagnosed Alzheimer’s disease GWAS and self-report proxy phenotype GWAX. Alzheimer’s Research & Therapy 14, 17 (2022).

22. European Alzheimer’s & Dementia Biobank Mendelian Randomization (EADB-MR) Collaboration, Genetic Associations Between Modifiable Risk Factors and Alzheimer Disease. JAMA Network Open 6, e2313734–e2313734 (2023).

23. J. G. Thorp et al., Genetic evidence that the causal association of educational attainment with reduced risk of Alzheimer’s disease is driven by intelligence. Neurobiology of Aging 119, 127–135 (2022).

24. Y. Chen et al., Genomic atlas of the plasma metabolome prioritizes metabolites implicated in human diseases. Nature Genetics 55, 44–53 (2023).

25. A. D. Grotzinger et al., Genomic structural equation modelling provides insights into the multivariate genetic architecture of complex traits. Nature Human Behaviour 3, 513–525 (2019).

26. Y. Wu, et al., Estimating genetic nurture with summary statistics of multigenerational genome-wide association studies. PNAS 118, e2023184118 (2021).

27. M. A. Nalls et al., Identification of novel risk loci, causal insights, and heritable risk for Parkinson’s disease: a meta-analysis of genome-wide association studies. The Lancet Neurology 18, 1091–1102 (2019).

28. W. van Rheenen et al., Genome-wide association analyses identify new risk variants and the genetic architecture of amyotrophic lateral sclerosis. Nature Genetics 48, 1043–1048 (2016).

29. R. Ferrari et al., Frontotemporal dementia and its subtypes: a genome-wide association study. The Lancet Neurology 13, 686–699 (2014).

30. R. Chia et al., Genome sequencing analysis identifies new loci associated with Lewy body dementia and provides insights into its genetic architecture. Nature Genetics 53, 294–303 (2021).

31. G. Mignogna et al., Patterns of item nonresponse behaviour to survey questionnaires are systematic and associated with genetic loci. Nature Human Behaviour, (2023).

32. T. Schoeler et al., Participation bias in the UK Biobank distorts genetic associations and downstream analyses. Nature Human Behaviour, (2023).

33. C. J. Willer, Y. Li, G. R. Abecasis, METAL: fast and efficient meta-analysis of genomewide association scans. Bioinformatics 26, 2190–2191 (2010).

34. J. C. Phelan, B. G. Link, in Medical Sociology on the Move. (Springer Netherlands, 2013), pp. 105–125.

35. J. M. Tublin, J. M. Adelstein, F. del Monte, C. K. Combs, L. E. Wold, Getting to the Heart of Alzheimer Disease. Circulation Research 124, 142–149 (2019).

36. D. A. Stakos et al., The Alzheimer’s Disease Amyloid-Beta Hypothesis in Cardiovascular Aging and Disease: JACC Focus Seminar. Journal of the American College of Cardiology 75, 952–967 (2020).

37. A. Fry et al., Comparison of sociodemographic and health-related characteristics of UK Biobank participants with those of the general population. American Journal of Epidemiology 186, 1026–1034 (2017).

38. N. Pirastu et al., Genetic analyses identify widespread sex-differential participation bias. Nature Genetics, (2021).

39. J. Tyrrell et al., Genetic predictors of participation in optional components of UK Biobank. Nature Communications 12, 886 (2021).

40. J. Mbatchou et al., Computationally efficient whole-genome regression for quantitative and binary traits. Nature Genetics, (2021).

41. P. Turley et al., Multi-trait analysis of genome-wide association summary statistics using MTAG. Nature Genetics 50, 229–237 (2018).

42. Q. Lu et al., A powerful approach to estimating annotation-stratified genetic covariance via GWAS summary statistics. Am J Hum Genet 101, 939–964 (2017).

43. B. Bulik-Sullivan et al., An atlas of genetic correlations across human diseases and traits. Nature Genetics 47, 1236–1241 (2015).

44. T. Ge, C.-Y. Chen, Y. Ni, Y.-C. A. Feng, J. W. Smoller, Polygenic prediction via Bayesian regression and continuous shrinkage priors. Nature Communications 10, 1776 (2019).

45. G. Hemani et al., The MR-Base platform supports systematic causal inference across the human phenome. eLife 7, e34408 (2018).

