## Supplementary Figures for "Pervasive biases in proxy GWAS based on parental history of Alzheimer’s disease"

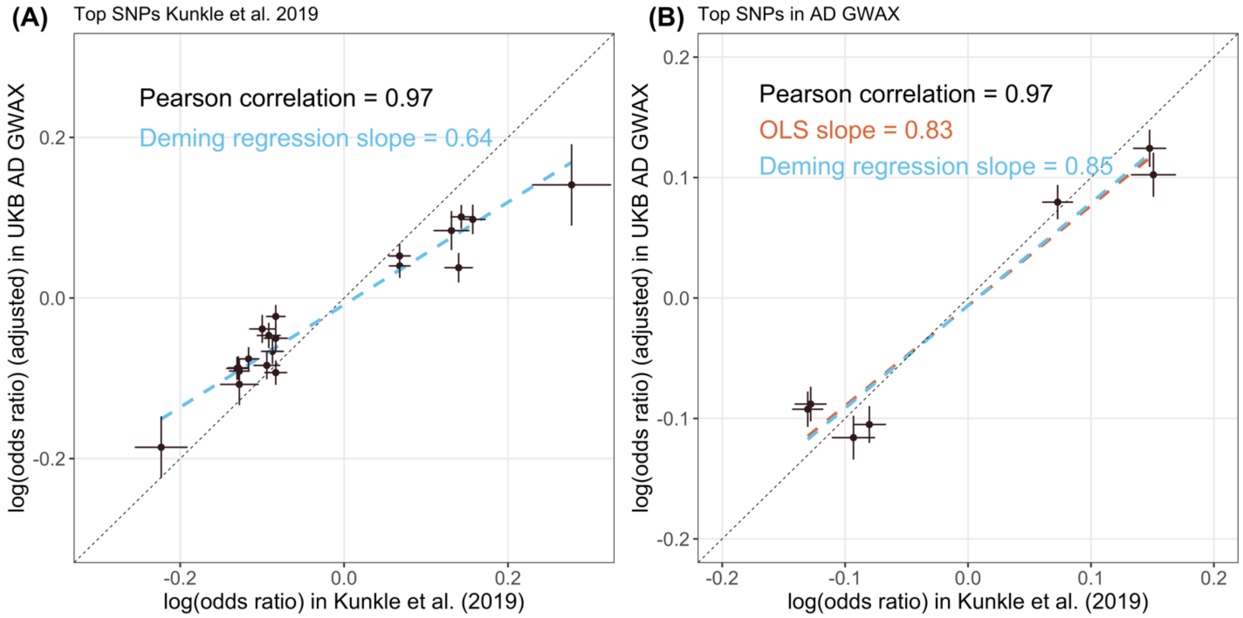


**Supplementary Figure 1. Comparing effect sizes (log odds ratio) in AD GWAX and GWAS (Kunkle et al. 2019).** (**A**) Genome-wide significant SNPs in Kunkle et al. 2019 GWAS and **(B)** AD GWAX. Dots and intervals denote point estimates and standard errors. The *APOE* variant (rs429358) is excluded due to its extreme effect size. We used Deming regression from the R package “mcr” to account for measurement error (**Methods**). Data for this plot can be found in **Supplementary Tables 1-2**.

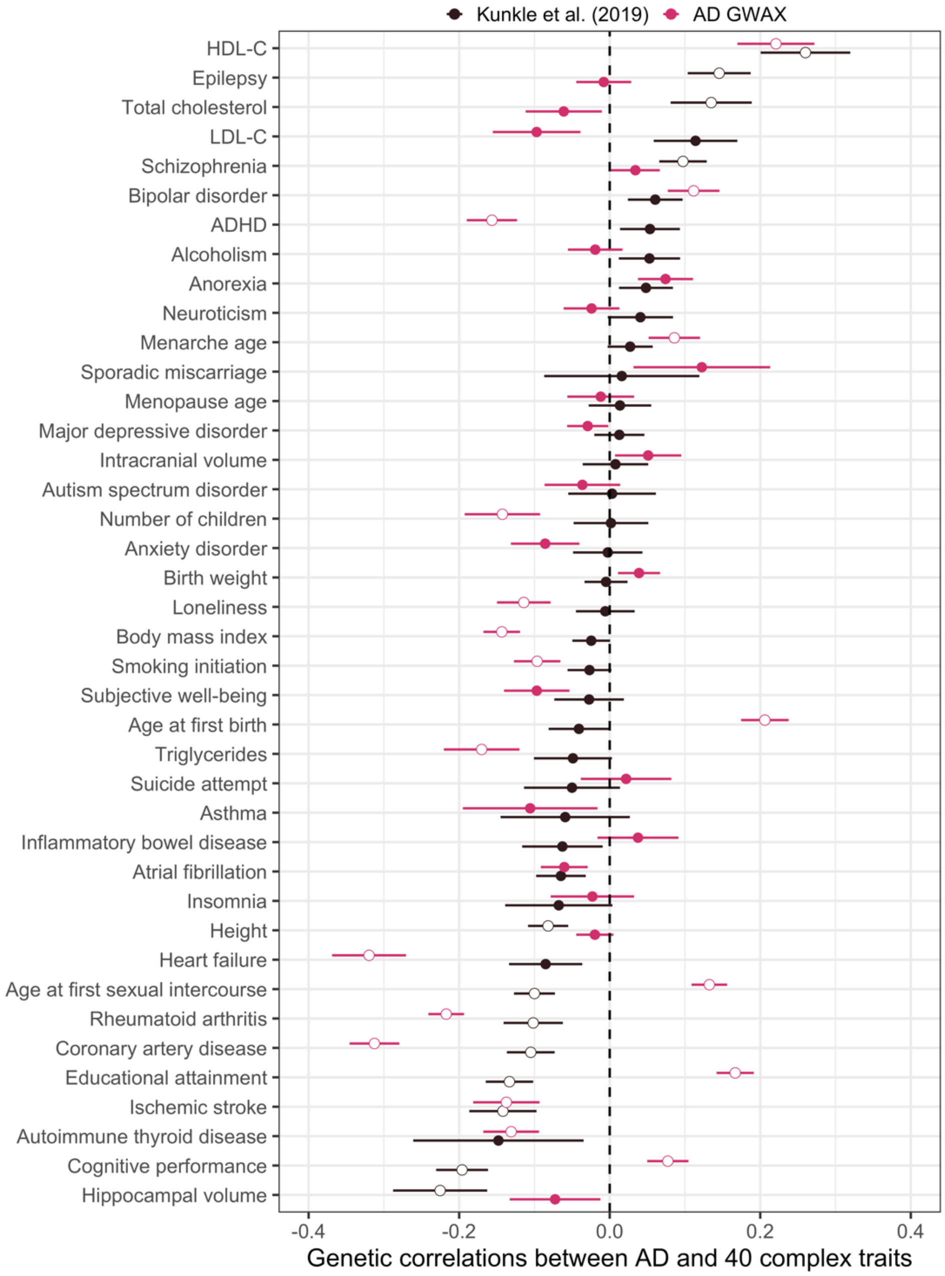


**Supplementary Figure 2. Genetic correlations between AD GWAX and GWAS (Kunkle et al. 2019) versus 40 complex traits.** Dots and intervals indicate point estimates and standard errors, respectively. Significant results at an FDR cutoff of 0.05 are highlighted with white circles. Traits were ordered based on genetic correlations with AD GWAS. HDL-C: high-density lipoprotein cholesterol, LDL-C: low-density lipoprotein cholesterol, ADHD: Attention-deficit/hyperactivity disorder. Data for this plot can be found in **Supplementary Tables 3-4**.

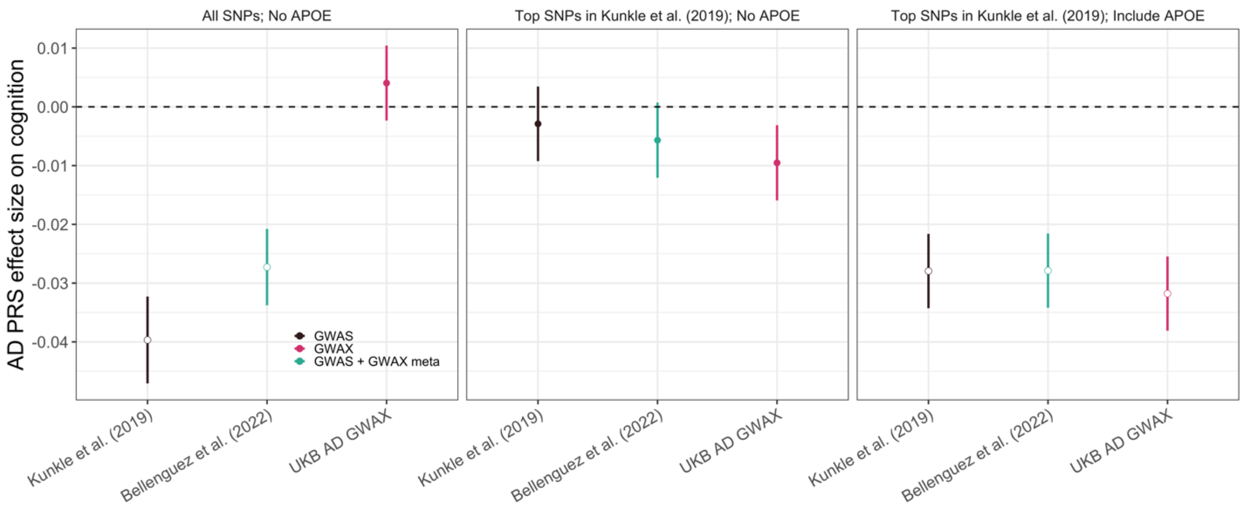


**Supplementary Figure 3. Association of AD PRS with late-life cognition in the HRS cohort.** PRS were computed using three different approaches (**Methods**): the left, middle, and right panels show results based on all variants except *APOE* (by PRS-CS), genome-wide significant loci identified in Kunkle et al. 2019, and genome-wide significant loci in Kunkle et al. 2019 with *APOE* excluded. Dots and intervals indicate the point estimates and +/- one standard error for the estimate, respectively. Significant results at an FDR cutoff of 0.05 are highlighted with white circles. Data for this plot can be found in **Supplementary Table 6**.

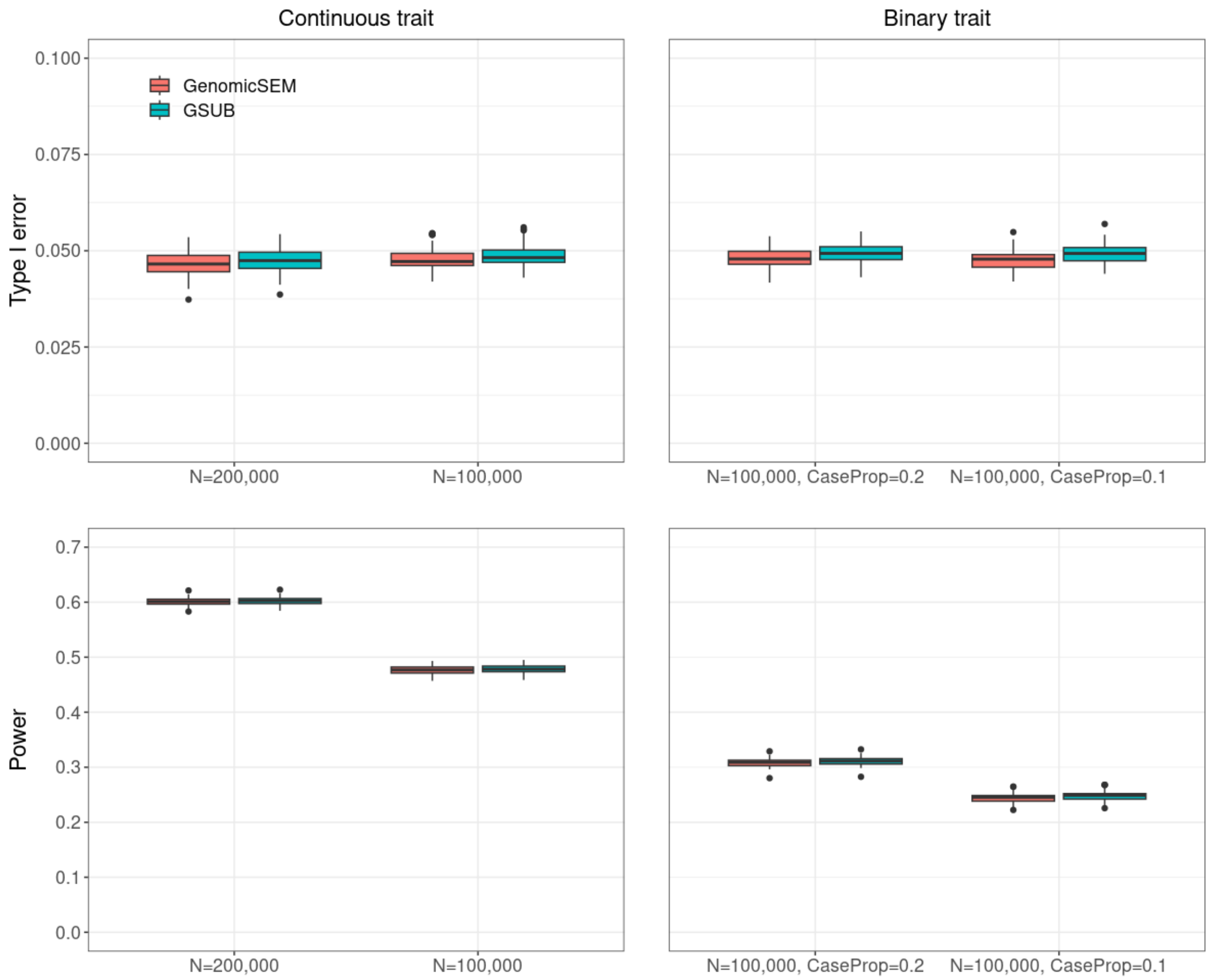


**Supplementary Figure 4. Comparison of GenomicSEM and GSUB for GWAS-by-subtraction.** We simulated both continuous and binary traits, then applied both the GenomicSEM and GSUB to conduct the GWAS-by-subtraction. We compared their type I error and power. We simulated both 100K and 200K sample sizes for the continuous traits. For the binary trait, we simulated 100K samples with case proportions of 0.1 and 0.2 (**Methods**).

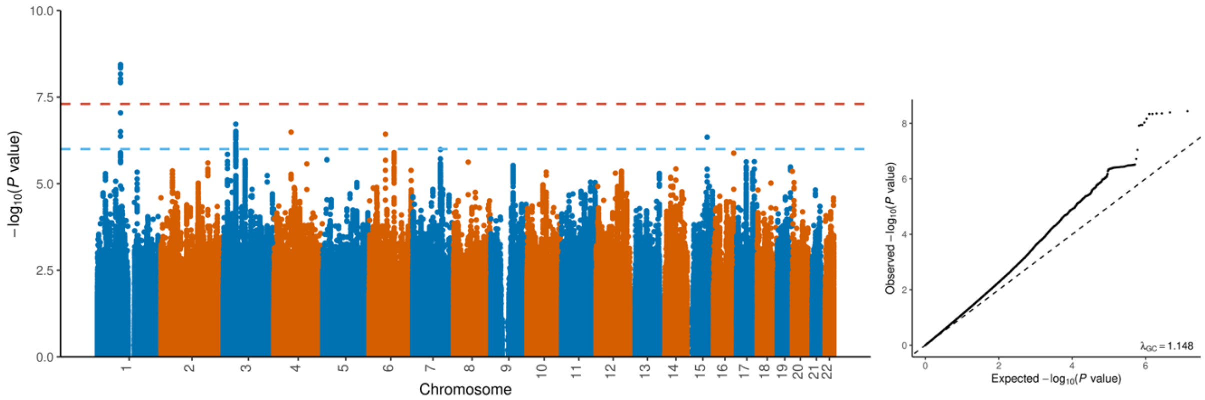


**Supplementary Figure 5. Manhattan and QQ plots for the GWAS on whether knowing the illnesses of parents in UKB**. N = 59,471 cases and 339,170 controls.

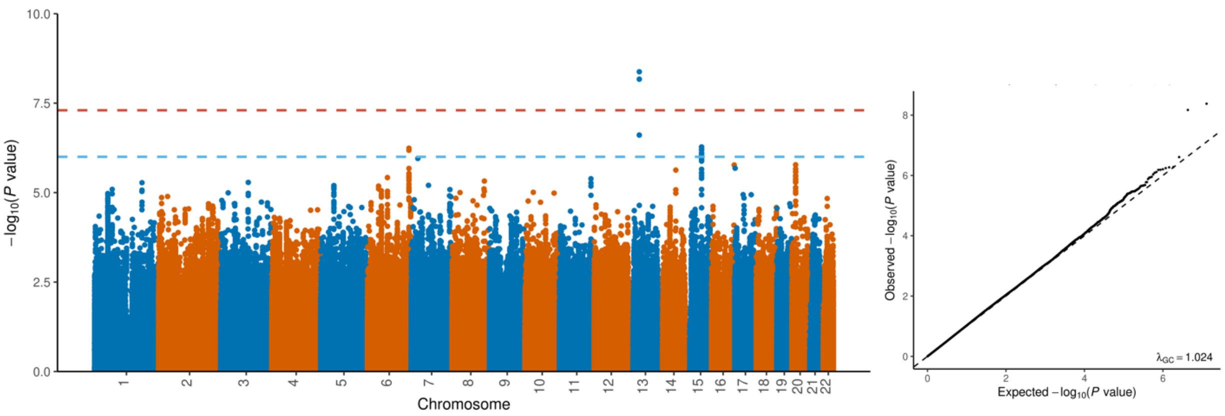


**Supplementary Figure 6. Manhattan and QQ plots for the GWAS on family medical history awareness in the AllofUs.** N = 77,579.

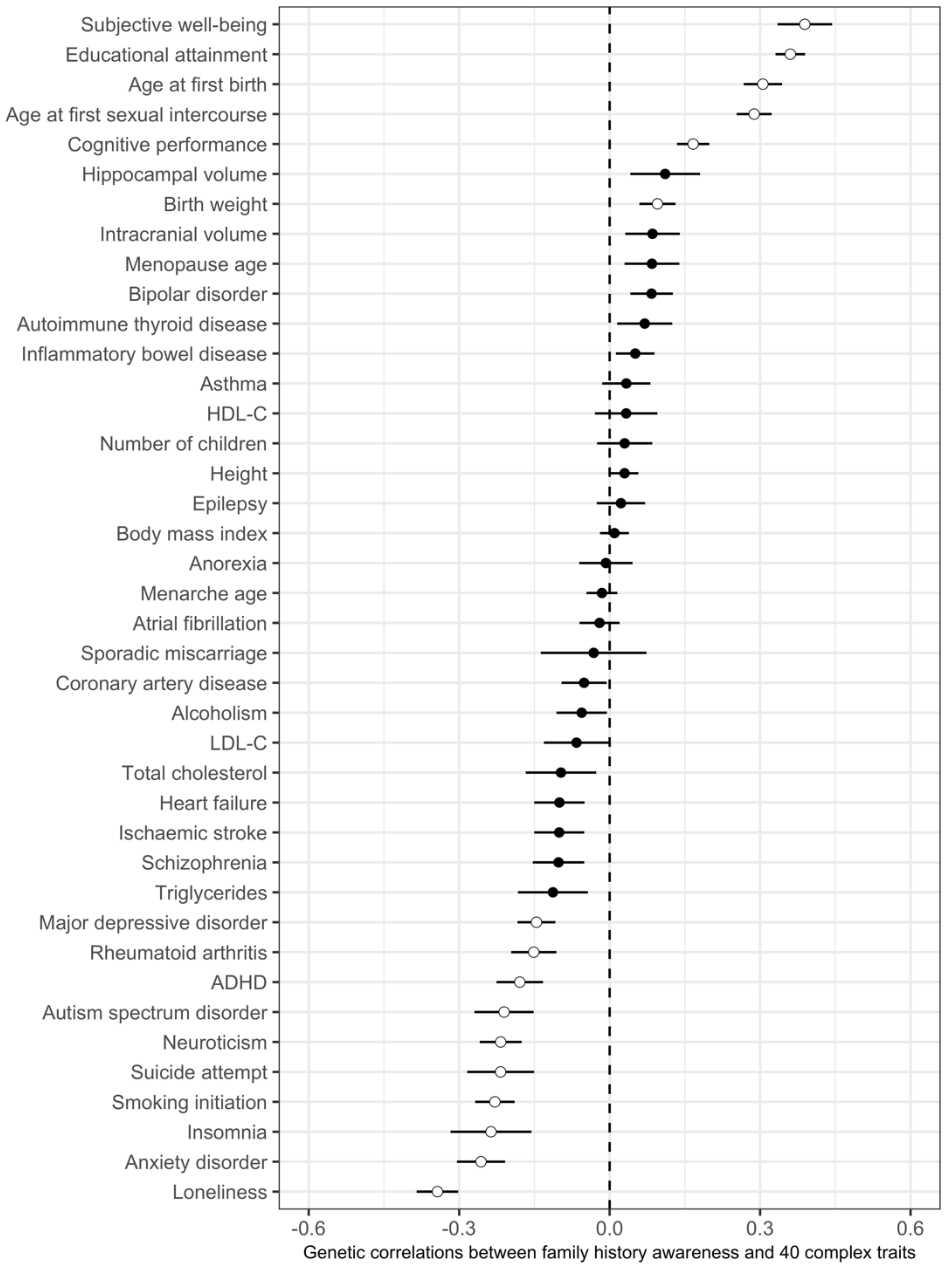


**Supplementary Figure 7. Genetic correlation between the awareness of family health history in AllofUs vs. 40 complex traits**. Dots and intervals indicate the point estimates and SE of genetic correlations, respectively. Significant correlations at a false discovery rate (FDR) cutoff of 0.05 are highlighted with white circles. Educational attainment NonCog: Non-cognitive performance component in educational attainment. Data for this plot can be found in the **Supplementary Table 12**.

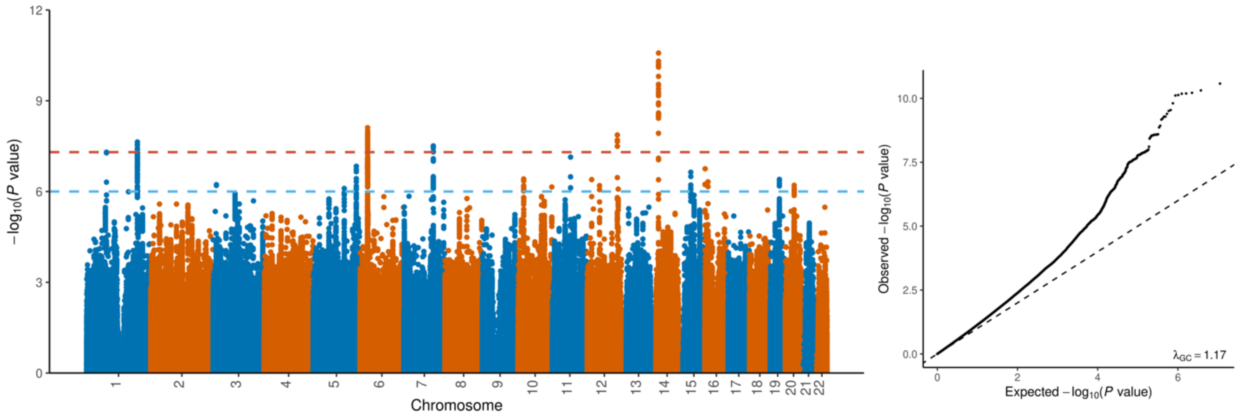


**Supplementary Figure 8. Manhattan and QQ plots for the GWAS on whether participating the personal and family health history survey in AllofUs**. N = 78,027 cases (participants) and 47,519 controls (non-participants).

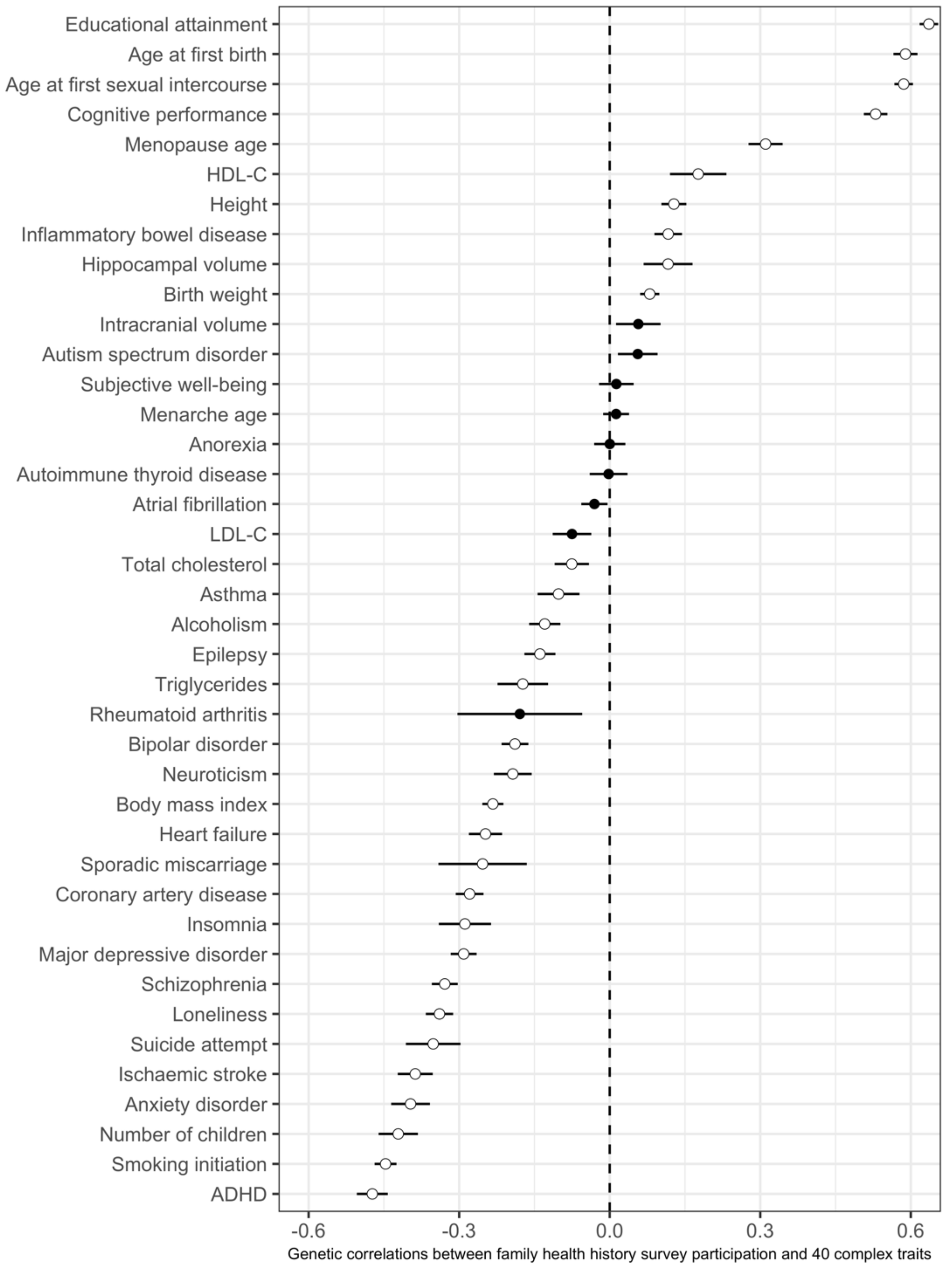


**Supplementary Figure 9. Genetic correlation between whether participating the optional personal and family health history survey in AllofUs vs. 40 complex traits.** Dots and intervals indicate the point estimates and SE of genetic correlations, respectively. Significant correlations at a false discovery rate (FDR) cutoff of 0.05 are highlighted with white circles. Data for this plot can be found in the **Supplementary Table 13**.

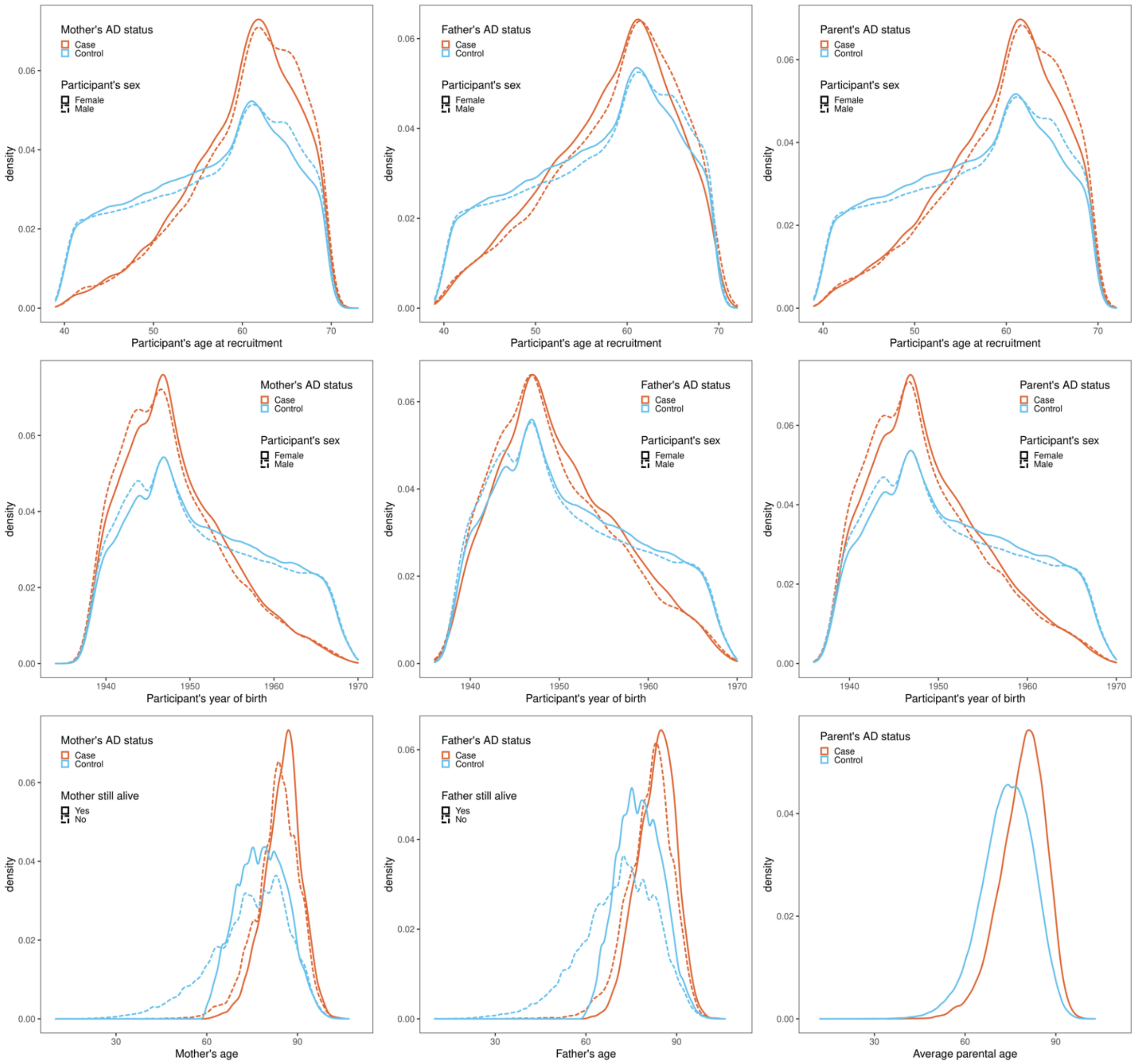


**Supplementary Figure 10. Comparing sample characteristics between AD proxy cases and proxy controls in UKB.** We compared the GWAX samples' age at recruitment, year of birth, and paternal/maternal/average-parental age.

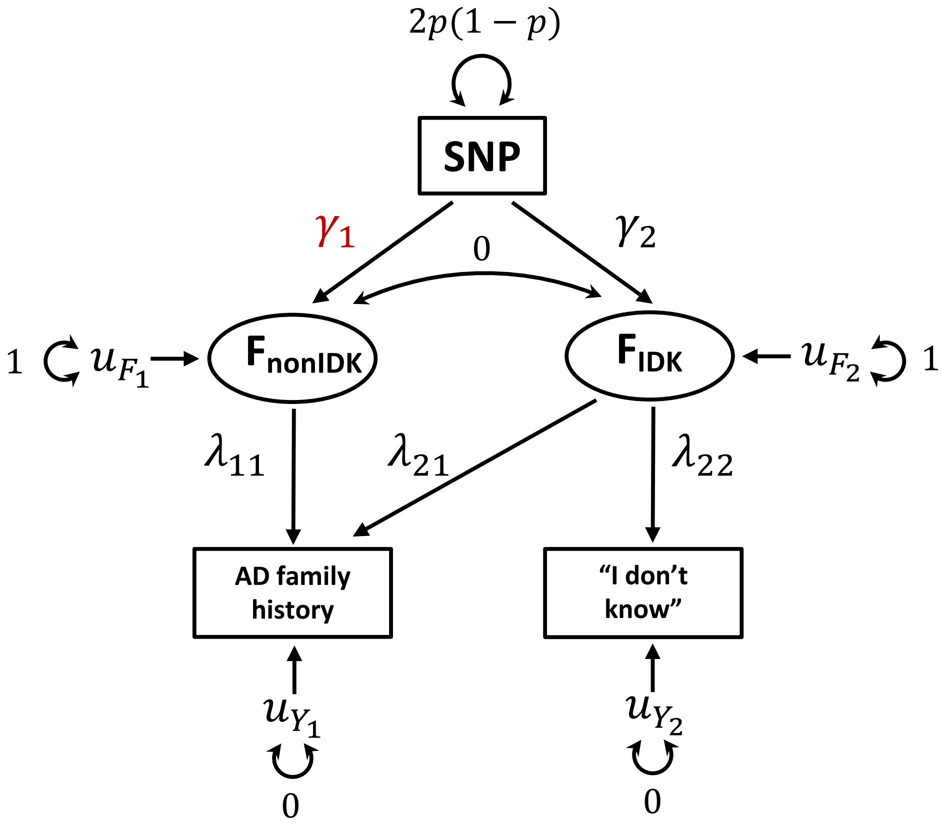


**Supplementary Figure 11. Schematic diagram for GWAS-by-subtraction.** The main goal is to estimate genetic associations $\gamma_1$ with the nonIDK factor $F_{nonIDK}$ underlying parental disease history after regressing out the "I don't know" (IDK) component $F_{IDK}$ from AD GWAX.

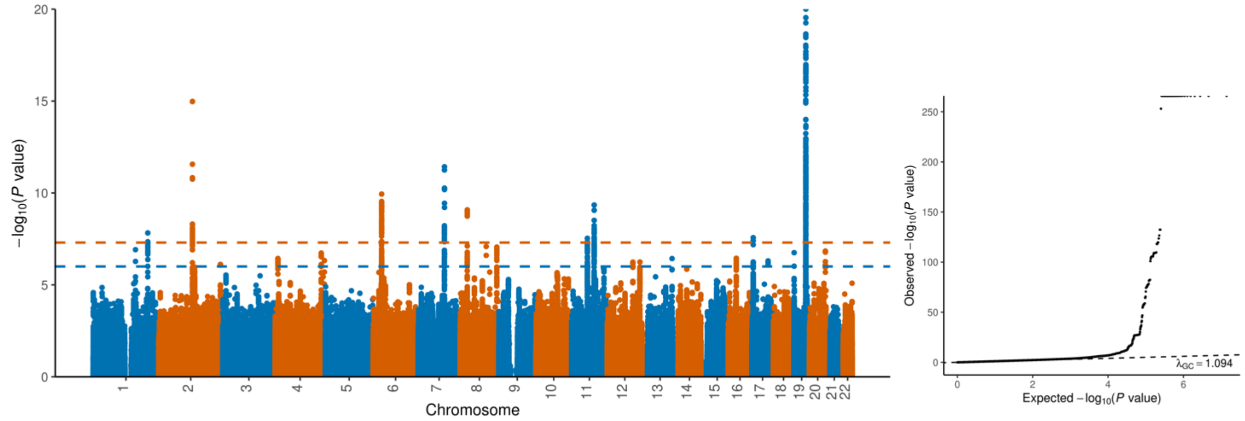


**Supplementary Figure 12. Manhattan and QQ plots of UKB AD GWAX where participants who selected "Do not know" were included in the controls.** We used only samples of European descent who are not adopted. N = 349,165 controls and 47,993 cases.

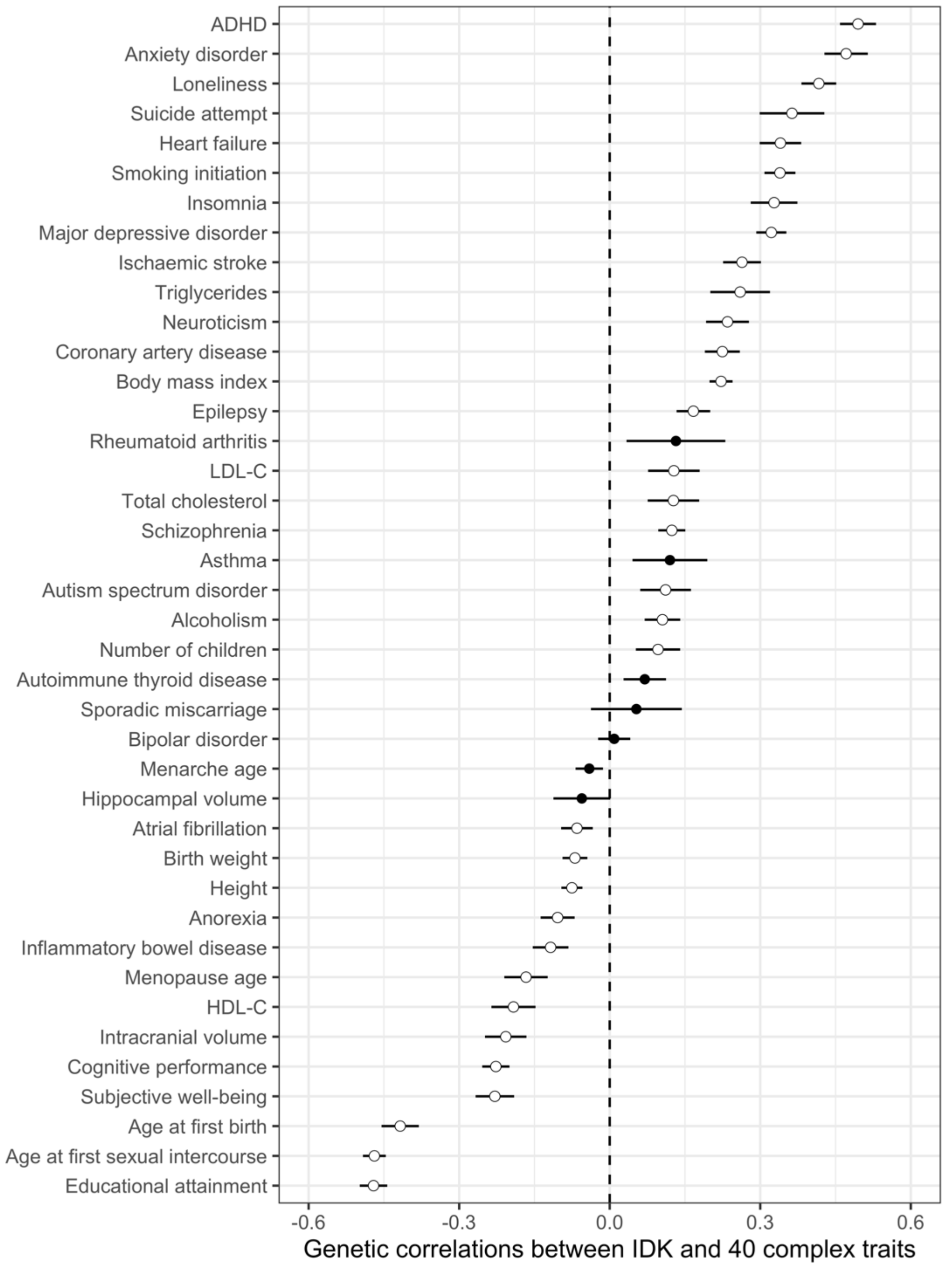


**Supplementary Figure 13. Genetic correlation between whether know the illness of parents vs. 40 complex traits.** Dots and intervals indicate the point estimates and SE of genetic correlations, respectively. Significant correlations at a false discovery rate (FDR) cutoff of 0.05 are highlighted with white circles. Data for this plot can be found in the **Supplementary Table 11**.

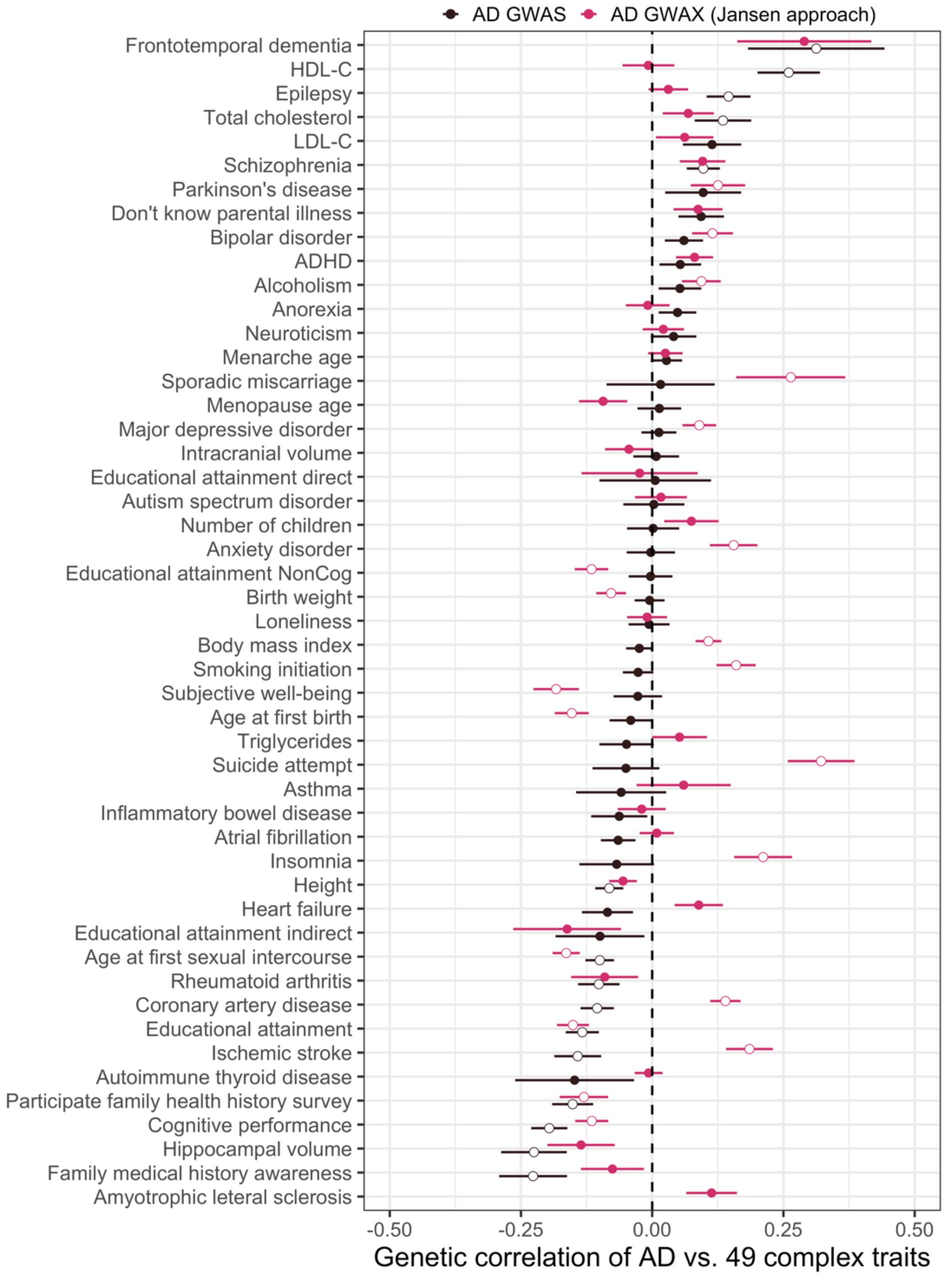


**Supplementary Figure 14. Genetic correlations of AD GWAS (Kunkle et al. 2019) and GWAX following Jansen et al. (2019) versus 49 complex traits.** Lewy body dementia was removed since it had NA results with both GWAS and GWAX. Dots and intervals indicate the point estimates and SE of genetic correlations, respectively. Significant correlations at a false discovery rate (FDR) cutoff of 0.05 are highlighted with white circles. Data for this plot can be found in the **Supplementary Table 16**.

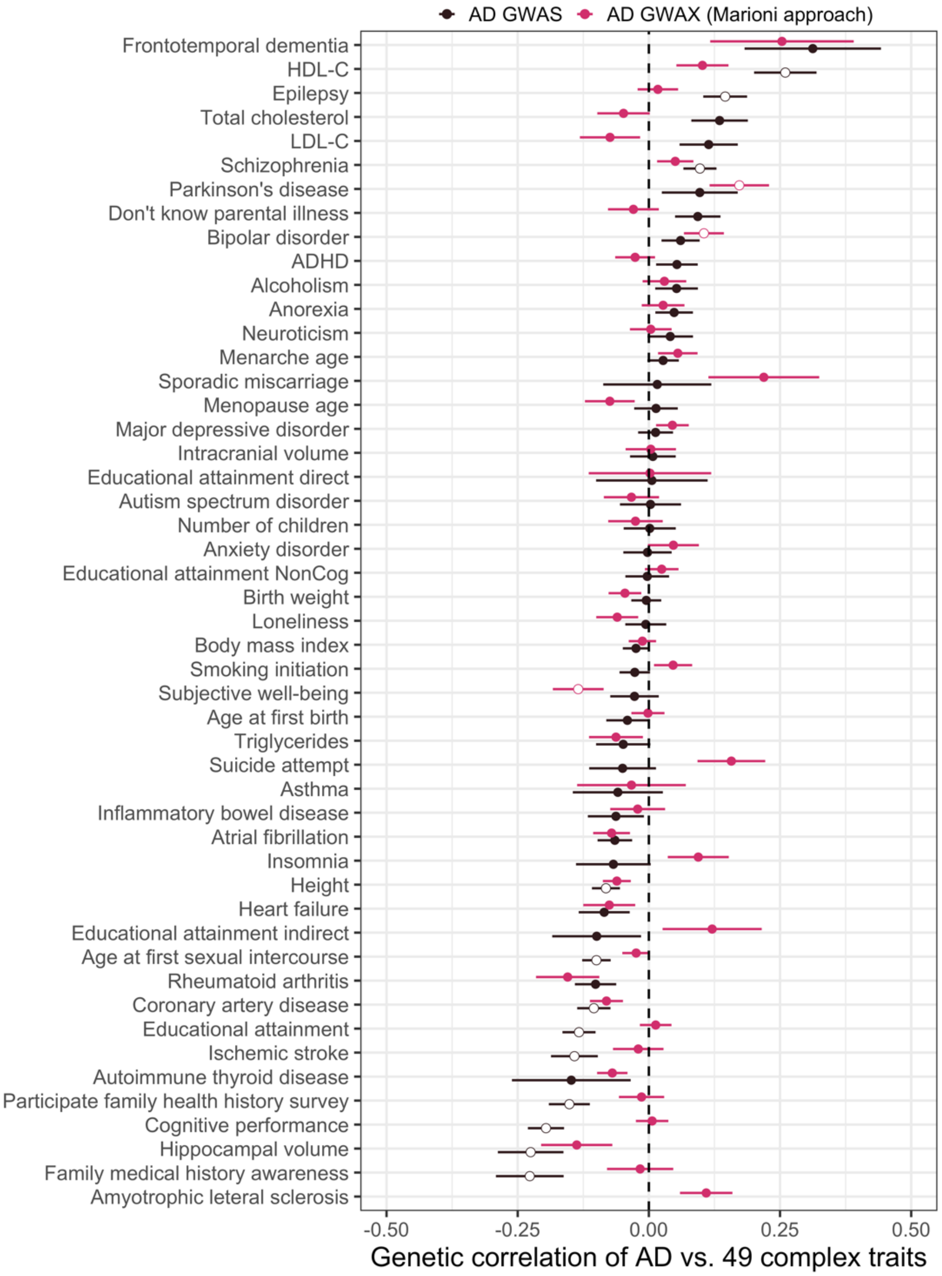


**Supplementary Figure 15. Genetic correlations of AD GWAS (Kunkle et al. 2019) and GWAX following Marioni et al. (2018) versus 49 complex traits.** Lewy body dementia was removed since it had NA results with both GWAS and GWAX. Dots and intervals indicate the point estimates and SE of genetic correlations, respectively. Significant correlations at a false discovery rate (FDR) cutoff of 0.05 are highlighted with white circles. Data for this plot can be found in the **Supplementary Table 17**.
